# Blinking fluorescent probes for tubulin nanoscopy in living and fixed cells

**DOI:** 10.1101/2021.06.01.446685

**Authors:** Rūta Gerasimaitė, Jonas Bucevičius, Kamila A. Kiszka, Georgij Kostiuk, Tanja Koenen, Gražvydas Lukinavičius

## Abstract

Here we report a small molecule probe for single molecule localisation microscopy (SMLM) of tubulin in living and fixed cells. We explored a series of constructs composed of taxanes and spontaneously blinking far-red dye hydroxymethyl silicon-rhodamine (HMSiR). We found that the linker length profoundly affects the probe permeability and off-targeting. The best performing probe, HMSiR-tubulin, is composed of cabazitaxel and 6’-regioisomer of HMSiR bridged by a C6 linker. Microtubule diameters of ≤50 nm can be routinely measured in SMLM experiments on living and fixed cells. HMSiR-tubulin also performs well in 3D stimulated emission depletion (STED) microscopy, allowing a complementary use of both nanoscopy methods for investigating microtubule functions in living cells.

---

As one of the major component of cytoskeleton, tubulin is involved in trafficking of biomolecules, cell movement and division. Paclitaxel is one of the most effective anti-cancer drugs used for the treatment of solid tumors such as ovarian, breast, and lung cancers^1^. It acts by stabilizing microtubules and thereby blocking cell progression through mitosis. It was originally isolated from the bark of the yew tree *Taxus brevifolia* in 1971^2^ and a tremendous repertoire of analogues has been reported by research groups and pharmaceutical companies ever since^1, 3^. Some of these analogues found their use in microscopy applications. Flutax-1 and Flutax-2 were the first fluorescent probes for tubulin imaging ^4–5^. They are composed of paclitaxel and fluorescein or Oregon Green, respectively. Further developments encompass various taxane analogs coupled to red-fluorescent dyes, such as boron-dipyrromethene (BODIPY) dyes, rhodamines, coumarines, carbo-, germano- or silicon-rhodamines ^4, 6–10^. All these probes are designed for the ensemble fluorescence microscopy methods. The development of tubulin probes for single molecule localization microscopy (SMLM) in living cells is however lagging behind, with only one published example composed of spiropyran derivative and colchicine, which requires UV illumination ^11^. We are bridging this gap by synthesizing taxane analogs coupled to spontaneously blinking fluorophore hydroxymethyl silicon-rhodamine (HMSiR)^12^.

Our previous work has repeatedly pointed out the importance of finding the optimal combination of a targeting moiety and a fluorophore^13–15^. Therefore, we explored several designs by coupling 6’-regioisomer of HMSiR to the often used analog docetaxel (DTX), or taxanes that are poorly recognized by multidrug resistance proteins – cabazitaxel (CTX) and larotaxel (LTX)^16–17^ (Figure 1a,b). The CTX-containing probe performed best and we fine-tuned its structure by varying a linker length and introducing 5’-regioisomer of HMSiR.

**Figure 1.**
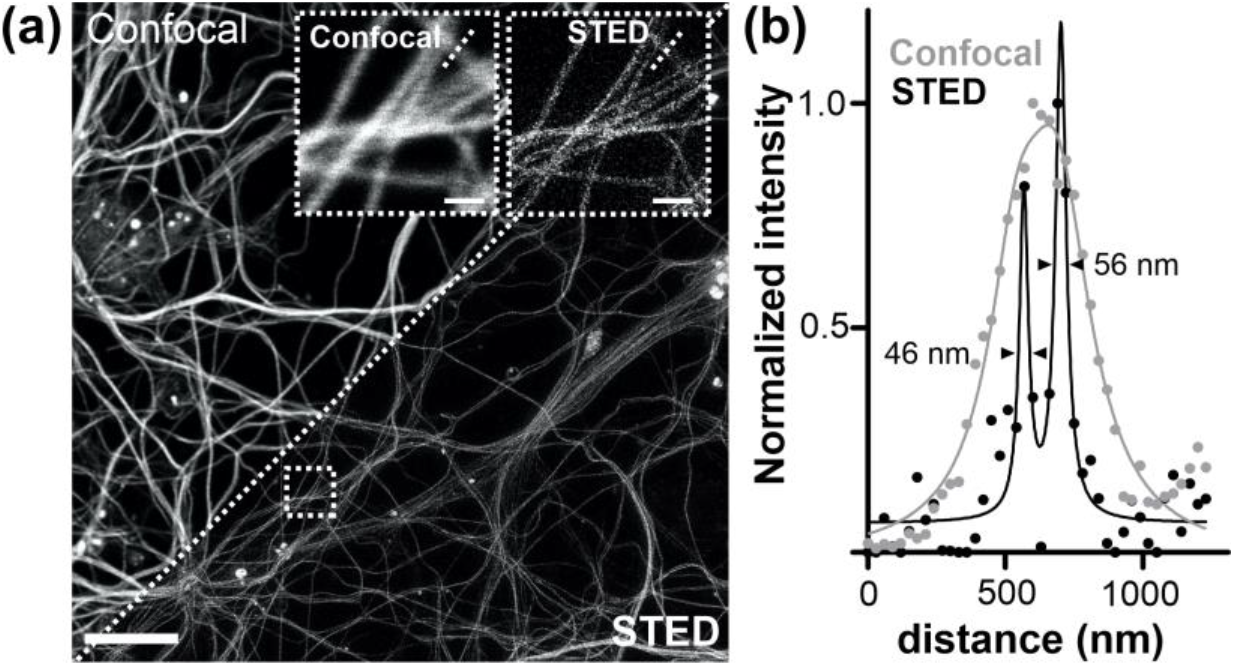
Tubulin probes synthesized in this study. (a) General structure of the probes, showing spirocyclization of the fluorophore that is responsible for spontaneous blinking. (b) Structure and naming convention of the probes. (c) Toxicity in Hela cells, determined as cell accumulation in subG1 cell cycle stage after 24 h incubation with the probes. Mean ± SD, N=3.

The 5-HMSiR-COOH and 6-HMSiR-COOH regioisomers were obtained by previously described procedures^12, 18^. The design of tubulin probes is based on the replacement of the *tert*-butyloxycarbonyl (Boc) group in taxanes with HMSiR tethered via linkers of variable length at the 3’-position (Figure 1a,b). This was achieved by performing peptide coupling reactions with 5/6-HMSIR-COOH dyes and taxanes bearing the linker with terminal amino group (Scheme S1) or by attaching required linkers with terminal carboxylic acid group to HMSiR moiety and further coupling it to N-*desboc*-taxane derivatives (Scheme S2).

To get an insight into the probes’ performance *in cellulo*, we stained U-2 OS cells and imaged them with a spinning disk confocal microscope. Microtubule staining by probes **5** and **6** was weak, largely overshadowed by bright staining of cytoplasmic vesicles (Figure S1a). In contrast, all CTX-based probes stained mainly microtubules, and probe **3** (HMSiR-tubulin) was the brightest. In none of the cases addition of verapamil^13–14^ had an effect, indicating that efflux pumps do not limit probe accumulation. Fixing cells with 0.2% glutaraldehyde followed by quenching with NaBH_4_ is known to preserve taxane interaction with tubulin^19–20^. All probes stained microtubules in fixed cells (Figure S1b), although the intensity varied. HMSiR-tubulin (**3**) and probe **1** were the brightest, indicating that the cell permeability of the latter probe is suboptimal. The remaining probes were dimmer, suggesting either reduced affinity and/or unfavourable interaction between fluorophore and tubulin. In fixed cells, probes **5** and **6** stained microtubules, indicating that the cell permeability is the major factor compromising their performance. In concordance, probes **5** and **6** showed low cytotoxicity, with half maximal inhibitory concentration (*IC_50_*) in the micromolar range (Figures 1c and S2). Consistent with the staining data, probe **1** was the least toxic and the least cell permeable in CTX miniseries. Importantly, toxicity does not limit the probes’ real life application, as it was measured during a prolonged incubation (24 h), which is not likely in a typical SMLM experiment.

At this point, we concluded that CTX-containing probes hold more promise for *in cellulo* tubulin imaging and focused further investigation on this series.

HMSiR dye exists in the equilibrium between a fluorescent and non-fluorescent/non-absorbing states (Figure 1a). This equilibrium is strongly shifted towards the dark spiroether state at physiological pH (~7.4). Only a small subset of molecules exist in a fluorescent state at a given time point leading to spontaneous appearance of spatially separated light emitters – blinking events^12^. This equilibrium is highly sensitive to the fluorophore environment, which can change upon target binding. In aqueous buffer (PBS), our probes showed nearly no fluorescence and displayed broad absorbance spectra dominated by light scattering, indicating aggregation (Figure 2a). Binding to tubulin resulted in appearance of fluorescence and absorbance peak with no signs of light scattering, indicating disassembly of the aggregates. The equilibrium shift upon tubulin binding results in 2 to 6-fold increase of absorbance and 10-40-fold increase of fluorescence (Figures 2b,c). The changes of absorbance are mainly determined by spirocyclization equilibrium, while fluorescence is additionally quenched by intramolecular interactions in the aggregates. We estimated that 1-3% of the probe molecules exist in the fluorescent form when bound to tubulin by comparing absorbance/fluorescence in the presence of tubulin to absorbance/fluorescence in ethanol + 0.1% TFA, which represents the maximum values (Figure 2d).

**Figure 2.**
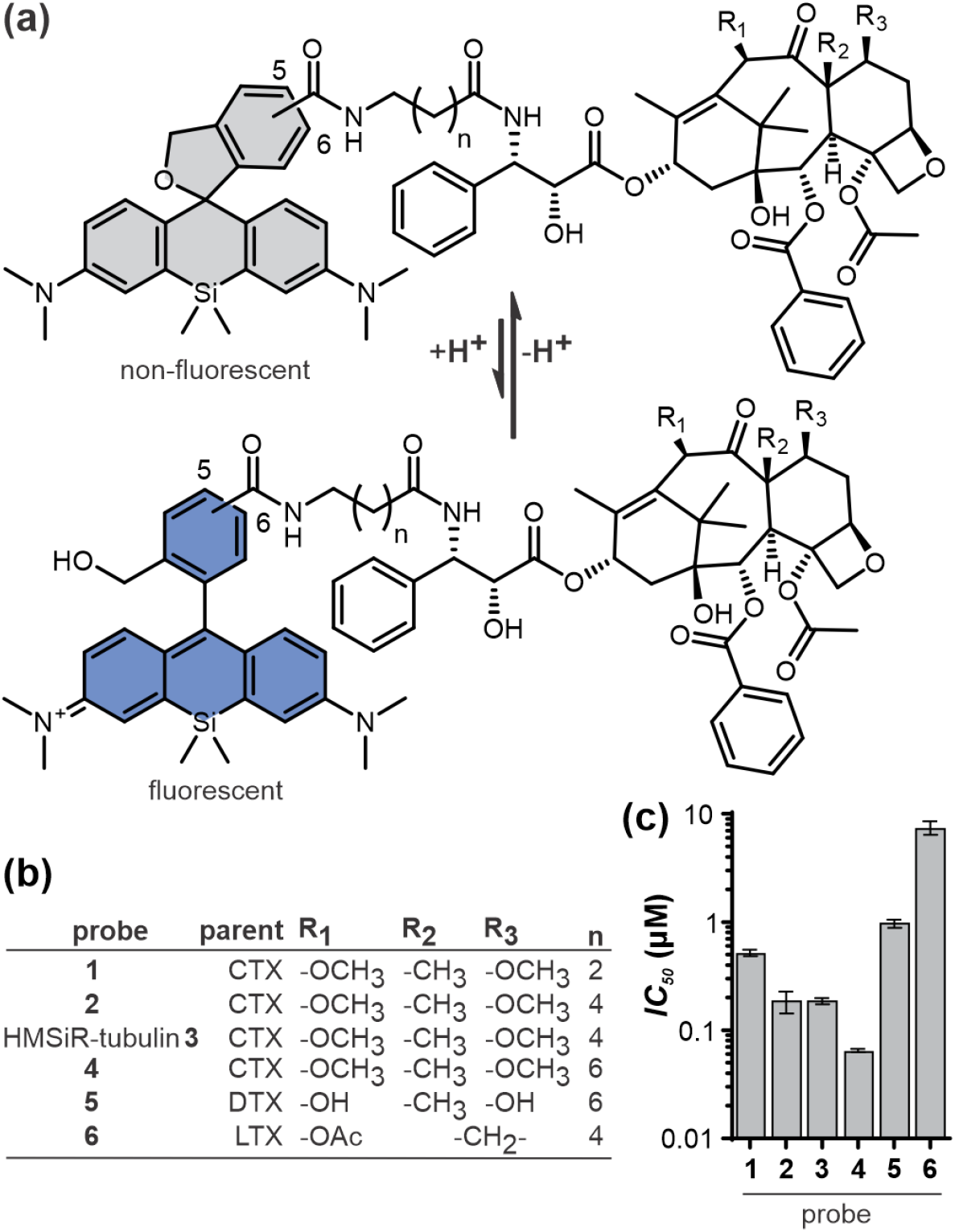
Properties of HMSiR CTX-based probes. (a) Normalized absorbance (solid line) and fluorescence (dashed line) spectra of **3**. PBS corr., absorbance spectrum corrected for light scattering. The inset zooms-in on weak signals in aqueous buffers. (b, c) Absorbance (659 nm) and fluorescence (673 nm) increase upon tubulin binding, as compared to PBS. (d) % of absorbing/fluo-rescent probe when bound to tubulin. Maximum values were determined in ethanol + 0.1% TFA. Mean ± SD, N=3 or 4. (e) apparent cyclization constants (pH at which half of molecules are in the spiroether state) of 5’- and 6’-regioisomers of free dye and HMSiR-CTX probes.

In addition to staining microtubules, HMSiR CTX-based probes can accumulate in intracellular vesicles, most likely acidic compartments. This was observed in several cell lines (Figures S1, S3) and in mouse primary hippocampal neurons (Figure S4). The trend was always the same: probe **4** showed the most, and HMSiR-tubulin (**3**) showed the least off-targeting. As weak bases, the rhodamine probes can be protonated and trapped in the acidic vesicle lumen. This can be rescued by increasing luminal pH, for example by applying milimolar concentration of NH_4_Cl. Indeed, adding 20 mM NH_4_Cl to human fibroblasts prestained with probes, substantially reduced the vesicle staining (Figure S3). Alternatively, low pH can shift the equilibrium towards the fluorescent state thus making the mistargeted probe more visible. We measured probe absorbance at different pH and found that apparent cyclisation constant (*pK_cyc_^app^*) of the probes is notably lower than that of the free dye^12^ and, likely, shifted by aggregation (Figures 2e and S5). However, as *pK_cyc_^app^* values were very close for all the probes, with that of probe **4** being the lowest (the most acidic), this mechanism cannot explain more prominent lysosome staining by this probe. Altogether, this data suggests that a long aliphatic linker favours probe mistargeting into acidic vesicles. Interestingly, despite poor staining of microtubules, probe **4** is substantially more toxic than others (Figure 1c), which might result from lysosome damaging.

HMSiR-tubulin (**3**) and probe **2** showed the best biocompatibility and staining. Although different only in fluorophore positional isomer, probe **2** consistently appeared less bright in living and (to a lesser degree) in fixed cells. We evaluated the binding affinity of these two probes by measuring the fluorescence intensity of the fixed cytoskeletons at variable probe concentrations. The binding curves could be fitted to a single-site dose response equation (Figure S6). The *K_D_^app^* for both probes were very similar and close to the values (10-100 nM) reported for Flutax probes^5, 20^. We found *K_D_^app^* = 121 ± 8 nM and 115 ± 8 nM, for HMSiR-tubulin (**3**) and probe **2**, respectively. Thus, differences in performance of these two probes do not result from different affinity to tubulin.

Next, we investigated the performance of HMSiR-tubulin (**3**) in SMLM imaging of microtubules in living and fixed cells. The movie of blinking fluorophores in living cells was acquired at 100 Hz frame rate with the laser power of 0.4 kW/cm^2^ in Hilo illumination mode. In the reconstructed images, the apparent diameter of the microtubule (FWHM) was 53 ± 13 nm (Figure 3a,e), which is lower than 79 nm and 69 nm reported for HMSiR- and HMCR550-labelled HaloTag-tubulin, respectively^12, 21^. The small label size and its binding inside the microtubule cylinder are the likely reasons for improved resolution. Under these conditions, we were able to observe microtubule dynamics (*i.e*. growing and shrinking) over 4 min (Supplementary Video 3). The resolution in living cells is limited by a low photon number detected per molecule per frame. Increasing the laser power led to increased photobleaching, while low frame rates resulted in too much motion blur. In fixed cells, imaging with 20 Hz increased the number of photons per molecule and a microtubule FWHM of 35 ± 6 nm was measured (Figure 3b, e), which is much closer to the one obtained by cryo-EM (~24 nm)^22^.

**Figure 3.**
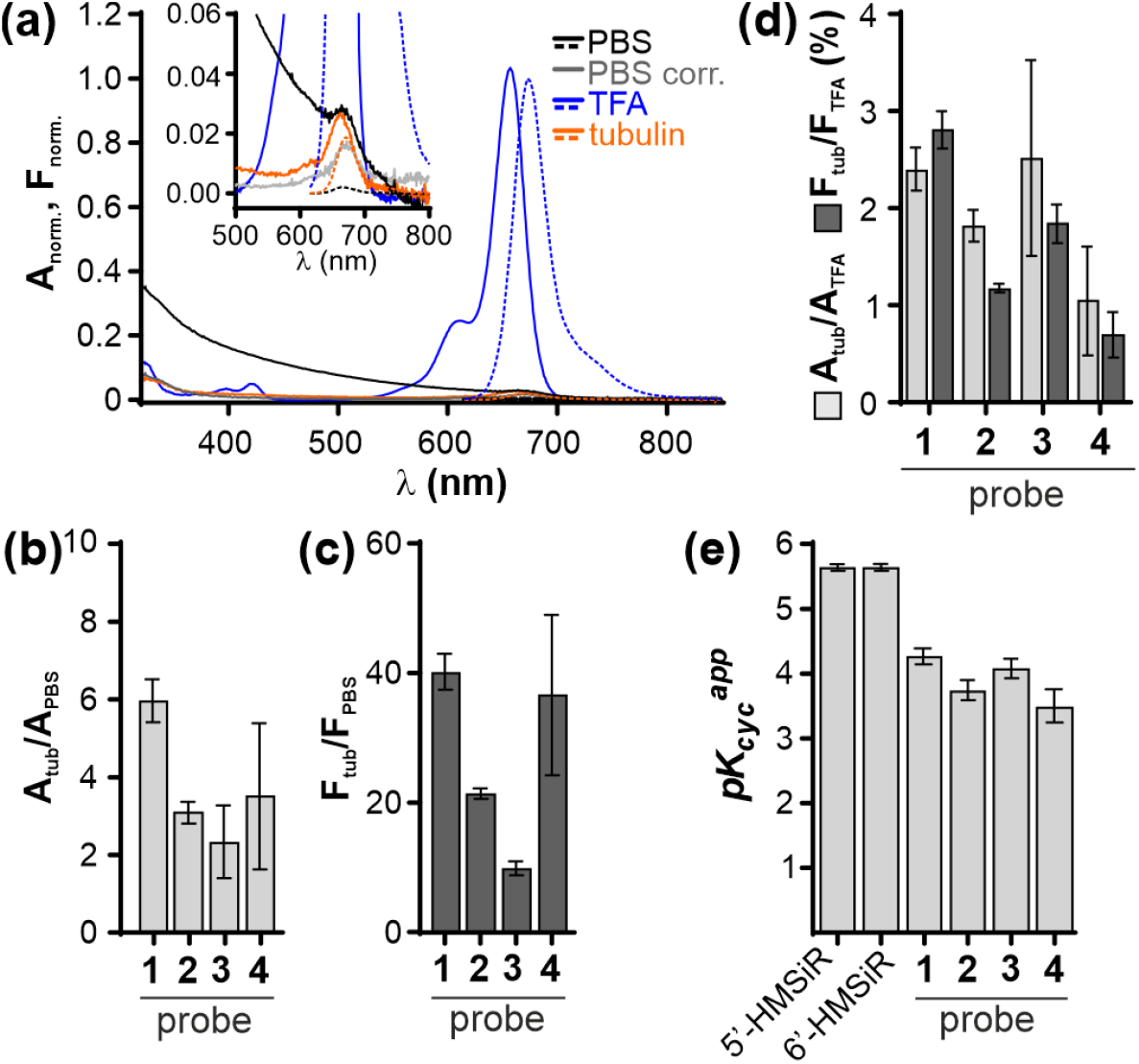
Imaging of microtubules in living and fixed U-2 OS cells stained with HMSiR-tubulin (**3**). (a) Living cells were stained with 100 nM **3** and imaged without washing at 100 Hz with 0.4 kW/cm^2^ 640 nm excitation power. The image was reconstructed from 2500 frames. (b) Fixed cells were stained with 10 nM **3** and imaged without washing at 20 Hz with 0.4 kW/cm^2^ 640 nm excitation power. The images were reconstructed from 4500 frames. Zoom-in on a single microtubule and intensity profile along the dashed line are shown, together with the corresponding fitting to the Gaussian function. Scale bar 2 μm, zoom-in – 0.5 μm. The movies used for reconstruction are in Supplementary Video 1 and 2. Fluorophore properties (histograms of localization uncertainty and photon count per molecule per frame and bleaching time-course) are shown. Fitting to lognormal distribution is shown as thick black line. Comparison of photon count per molecule per frame (c), localization uncertainty (d) and averaged microtubule FWHM between living and fixed cells (e). The statistics was obtained from 6 and 7 fields of view, from 2 independently prepared samples.

Both in living and in fixed cells, we saw little fluorescence recovery when prolonged illumination was alternated with incubation in the dark (Figure S7a,b). This has been observed before with a non-blinking probe SiR-tubulin^23^ and can be explained by a high (millimolar) local concentration of taxane binding sites on the microtubule. Consistently with slow exchange, we were able to image fixed microtubules several hours after washing away the probe.

A probe with a more prevalent dark state can be advantageous for imaging densely labelled structures. Therefore we examined probe **2** in the SMLM experiment. In fixed cells, it yielded images of similar quality as **3** (Figure S8). However, dimmer staining made the choice of imaging region in living cells extremely difficult. In summary, probe **3**, HMSiR-tubulin (**3**), showed the best staining and biocompatibility, and is the probe of choice for SMLM of tubulin.

In addition, we found that HMSiR-tubulin can be used for 2D and 3D STED nanoscopy of densely packed tubulin cables in living neurons, although the brightness is low and high excitation laser power is required (Figure 4 and Supplementary Video 4). Similar to the previously reported DNA probes^18^, we did not observe blinking. FWHM of a microtubule determined by 2D STED was similar to SMLM (~50 nm) (Figure 4b). In z-axis, microtubule FWHM of 300 nm was obtained, an ~2.5-fold improvement over standard confocal resolution (~700 nm) (Figure S9).

**Figure 4.**
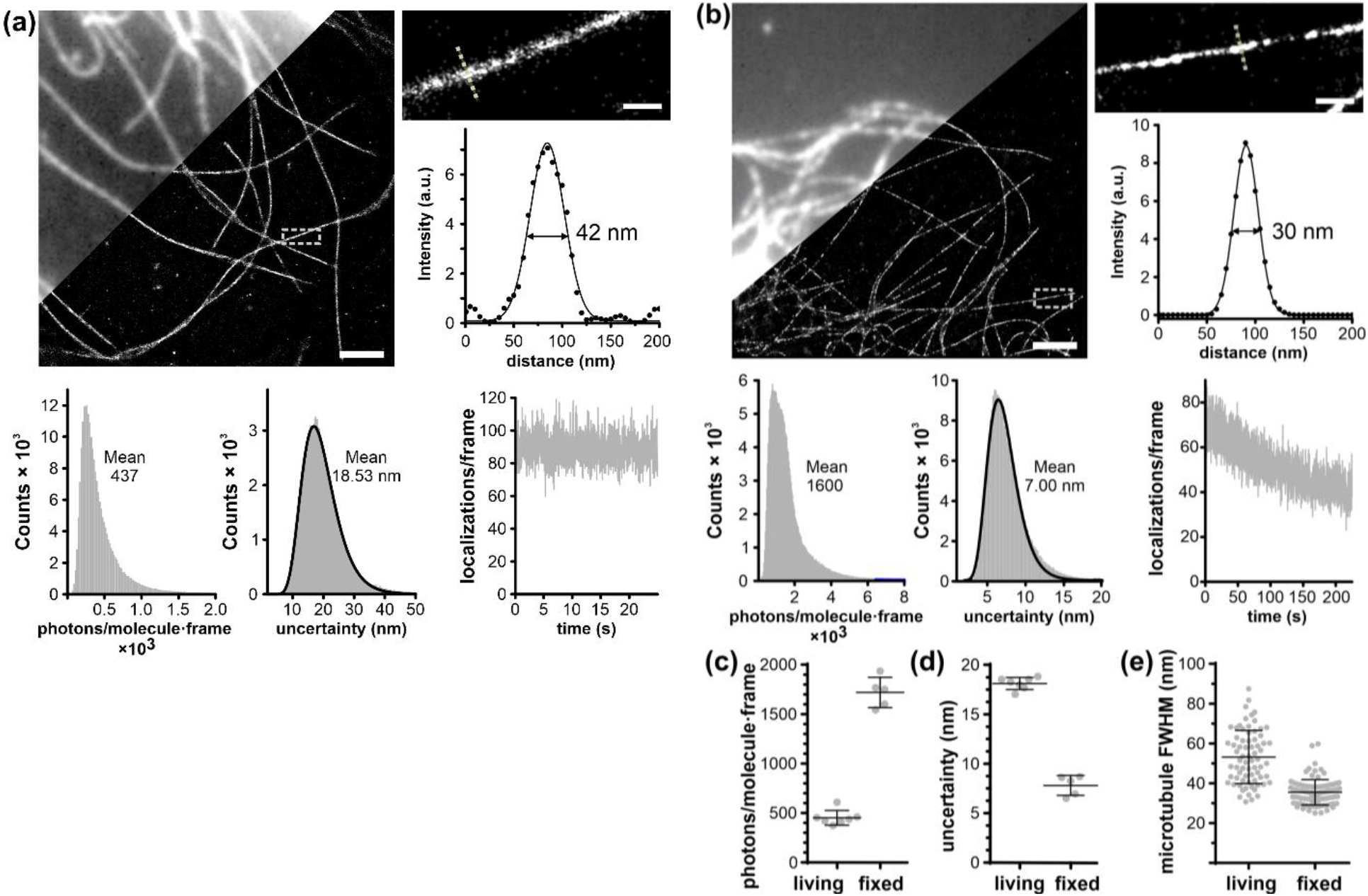
2D STED nanoscopy image of living mouse neurons stained with HMSiR-tubulin **3**. (a) Comparison of confocal and 2D STED with 775 nm images. The cells were stained with 300 nM HMSiR-tubulin for 1 h in growth medium and imaged without probe removal. Inset: zoom-in image of the region marked with dashed square. Scale bar 10 μm, inset – 1 μm. (b) Line profile of the region marked with dashed line in (a). Numbers represent FWHM.

In conclusion, we employed ligand and linker optimization to obtain HMSiR-tubulin, the first spontaneously blinking cell permeable probe for SMLM of microtubules in living and fixed cells. Furthermore, it is also compatible with STED using 775 nm depletion laser, thereby allowing interchangeable use of both techniques for investigating microtubule-related processes in living cells. Microtubule FWHM of 50 nm is reproducibly achieved by both methods while imaging dynamic tubulin structures.

## ASSOCIATED CONTENT

### Supporting Information

Supporting Information is available free of charge on the ACS Publications website.

Supplementary figures, materials and methods, synthetic procedures and characterization of the compounds (PDF).

Supplementary Video 1, blinking of HMSiR-tubulin in living U-2 OS cells.

Supplementary Video 2, blinking of HMSiR-tubulin in fixed U-2 OS cells

Supplementary Video 3, tubulin dynamics in living U-2 OS cells over 4 min.

Supplementary Video 4, 3D STED imaging of tubulin in primary mouse neurons.

## AUTHOR INFORMATION

### Author Contributions

G.L. conceived the project; R.G., J.B., K.K., G.K., T.G. and G.L. performed the experiments and analyzed the data. R.G., J.B. and G.L. wrote the initial draft; all authors contributed to the final version of the manuscript.

### Notes

The authors declare no competing financial interests.

## ACKNOWLEDGMENT

The authors acknowledge funding by the Max Planck Society. J.B. is grateful to the Max Planck Society for a Nobel Laureate Fellowship. G.K. is a recipient of EMBO Long Term Fellowship (ALTF 135-2019). The authors are grateful to Dr. Vladimir Belov, Jan Seikowski, Jens Schimpfhauser and Jürgen Bienert for the NMR/ESI-MS measurements of numerous compounds. We are grateful to dr. Peter Lenart and dr. Antonio Politi for their help with SMLM imaging.

## REFERENCES

1. Rohena, C. C.; Mooberry, S. L., Recent progress with microtubule stabilizers: new compounds, binding modes and cellular activities. Nat. Prod. Rep. 2014, 31 (3), 335–55.

2. Wani, M. C.; Taylor, H. L.; Wall, M. E.; Coggon, P.; McPhail, A. T., Plant antitumor agents. VI. The isolation and structure of taxol, a novel antileukemic and antitumor agent from Taxus brevifolia. J. Am. Chem. Soc. 1971, 93 (9), 2325–7.

3. Wang, Y. F.; Shi, Q. W.; Dong, M.; Kiyota, H.; Gu, Y. C.; Cong, B., Natural taxanes: developments since 1828. Chem. Rev. 2011, 111 (12), 7652–709.

4. Barasoain, I.; Diaz, J. F.; Andreu, J. M., Fluorescent taxoid probes for microtubule research. Methods Cell Biol. 2010, 95, 353–72.

5. Diaz, J. F.; Strobe, R.; Engelborghs, Y.; Souto, A. A.; Andreu, J. M., Molecular recognition of taxol by microtubules. Kinetics and thermodynamics of binding of fluorescent taxol derivatives to an exposed site. J. Biol. Chem. 2000, 275 (34), 26265–76.

6. Lukinavičius, G.; Reymond, L.; D’Este, E.; Masharina, A.; Gottfert, F.; Ta, H.; Guther, A.; Fournier, M.; Rizzo, S.; Waldmann, H.; Blaukopf, C.; Sommer, C.; Gerlich, D. W.; Arndt, H. D.; Hell, S. W.; Johnsson, K., Fluorogenic probes for live-cell imaging of the cytoskeleton. Nat. Methods 2014, 11 (7), 731–3.

7. Evangelio, J. A.; Abal, M.; Barasoain, I.; Souto, A. A.; Lillo, M. P.; Acuna, A. U.; Amat-Guerri, F.; Andreu, J. M., Fluorescent taxoids as probes of the microtubule cytoskeleton. Cell Motil. Cytoskeleton 1998, 39 (1), 73–90.

8. Butkevich, A. N.; Ta, H.; Ratz, M.; Stoldt, S.; Jakobs, S.; Belov, V. N.; Hell, S. W., Two-Color 810 nm STED Nanoscopy of Living Cells with Endogenous SNAP-Tagged Fusion Proteins. ACS Chem. Biol. 2017.

9. Butkevich, A. N.; Belov, V. N.; Kolmakov, K.; Sokolov, V. V.; Shojaei, H.; Sidenstein, S. C.; Kamin, D.; Matthias, J.; Vlijm, R.; Engelhardt, J.; Hell, S. W., Hydroxylated Fluorescent Dyes for Live-Cell Labeling: Synthesis, Spectra and Super-Resolution STED. Chemistry – A European Journal 2017, 23 (50), 12114–12119.

10. Lee, M. M.; Gao, Z.; Peterson, B. R., Synthesis of a Fluorescent Analogue of Paclitaxel That Selectively Binds Microtubules and Sensitively Detects Efflux by P-Glycoprotein. Angew. Chem. Int. Ed. Engl. 2017, 56 (24), 6927–6931.

11. Zhang, H.; Wang, C.; Jiang, T.; Guo, H.; Wang, G.; Cai, X.; Yang, L.; Zhang, Y.; Yu, H.; Wang, H.; Jiang, K., Microtubule-targetable fluorescent probe: site-specific detection and super-resolution imaging of ultratrace tubulin in microtubules of living cancer cells. Anal Chem 2015, 87 (10), 5216–22.

12. Uno, S. N.; Kamiya, M.; Yoshihara, T.; Sugawara, K.; Okabe, K.; Tarhan, M. C.; Fujita, H.; Funatsu, T.; Okada, Y.; Tobita, S.; Urano, Y., A spontaneously blinking fluorophore based on intramolecular spirocyclization for live-cell super-resolution imaging. Nat Chem 2014, 6 (8), 681–9.

13. Bucevičius, J.; Keller-Findeisen, J.; Gilat, T.; Hell, S. W.; Lukinavičius, G., Rhodamine-Hoechst positional isomers for highly efficient staining of heterochromatin. Chem Sci 2019, 10 (7), 1962–1970.

14. Gerasimaitė, R.; Seikowski, J.; Schimpfhauser, J.; Kostiuk, G.; Gilat, T.; D’Este, E.; Schnorrenberg, S.; Lukinavičius, G., Efflux pump insensitive rhodamine-jasplakinolide conjugates for G-and F-actin imaging in living cells. Org Biomol Chem 2020, 18 (15), 2929–2937.

15. Lukinavičius, G.; Mitronova, G. Y.; Schnorrenberg, S.; Butkevich, A. N.; Barthel, H.; Belov, V. N.; Hell, S. W., Fluorescent dyes and probes for super-resolution microscopy of microtubules and tracheoles in living cells and tissues. Chem Sci 2018, 9 (13), 3324–3334.

16. Abidi, A., Cabazitaxel: A novel taxane for metastatic castration-resistant prostate cancer-current implications and future prospects. J. Pharmacol. Pharmacother. 2013, 4 (4), 230–7.

17. Metzger-Filho, O.; Moulin, C.; de Azambuja, E.; Ahmad, A., Larotaxel: broadening the road with new taxanes. Expert Opin. Investig. Drugs. 2009, 18 (8), 1183–9.

18. Bucevičius, J.; Gilat, T.; Lukinavičius, G., Far-red switching DNA probes for live cell nanoscopy. Chem Commun (Camb) 2020, 56 (94), 14797–14800.

19. Barasoain, I.; Diaz, J. F.; Andreu, J. M., Fluorescent taxoid probes for microtubule research. Methods Cell Biol 2010, 95, 353–72.

20. Diaz, J. F.; Barasoain, I.; Andreu, J. M., Fast kinetics of Taxol binding to microtubules. Effects of solution variables and microtubule-associated proteins. J Biol Chem 2003, 278 (10), 8407–19.

21. Tachibana, R.; Kamiya, M.; Morozumi, A.; Miyazaki, Y.; Fujioka, H.; Nanjo, A.; Kojima, R.; Komatsu, T.; Ueno, T.; Hanaoka, K.; Yoshihara, T.; Tobita, S.; Urano, Y., Design of spontaneously blinking fluorophores for live-cell super-resolution imaging based on quantum-chemical calculations. Chem Commun (Camb) 2020, 56 (86), 13173–13176.

22. Kellogg, E. H.; Hejab, N. M. A.; Howes, S.; Northcote, P.; Miller, J. H.; Diaz, J. F.; Downing, K. H.; Nogales, E., Insights into the Distinct Mechanisms of Action of Taxane and Non-Taxane Microtubule Stabilizers from Cryo-EM Structures. J. Mol. Biol. 2017, 429 (5), 633–646.

23. Spahn, C.; Grimm, J. B.; Lavis, L. D.; Lampe, M.; Heilemann, M., Whole-Cell, 3D, and Multicolor STED Imaging with Exchangeable Fluorophores. Nano Lett 2019, 19 (1), 500–505.

