## Supplementary material for "Blinking fluorescent probes for tubulin nanoscopy in living and fixed cells" (from another version of this preprint: version 1, 2021-06-02)

### Supplementary figures

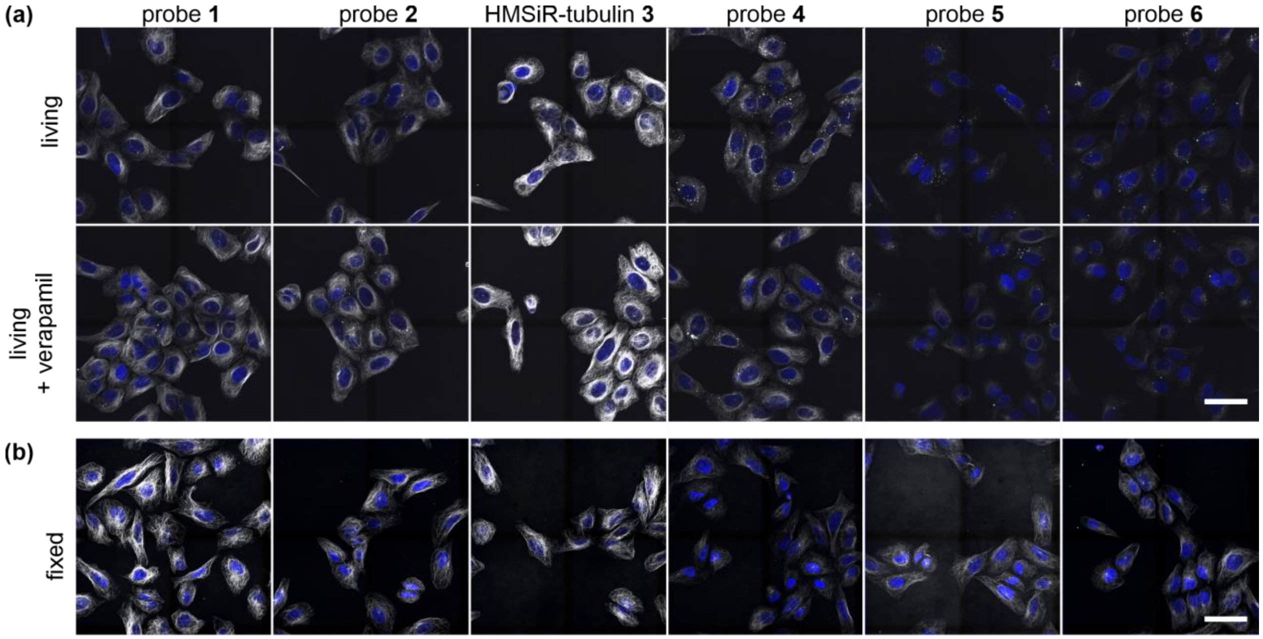

**Figure S1. Staining U-2 OS cells with tubulin probes.**

**(a)** Living U-2 OS cells were incubated with 300 nM probe, +/-10 µM verapamil and 1 µg/ml Hoechst 33342 in DMEM medium with FBS for 1h at 37°C and imaged on confocal spinning disk microscope without washing. **(b)** The cells were fixed as described in Methods section and incubated with 100 nM probe and 0.1 µg/ml Hoechst 33342 in PEM buffer for 30 min. at room temperature. In all cases, maximal intensity projections of 39 planes (living cells) or 41 planes (fixed cells), acquired with a step size of 200 nm, are shown. Probe channel is greyscale, Hoechst 33342 is blue. Scale bar – 50 µm.

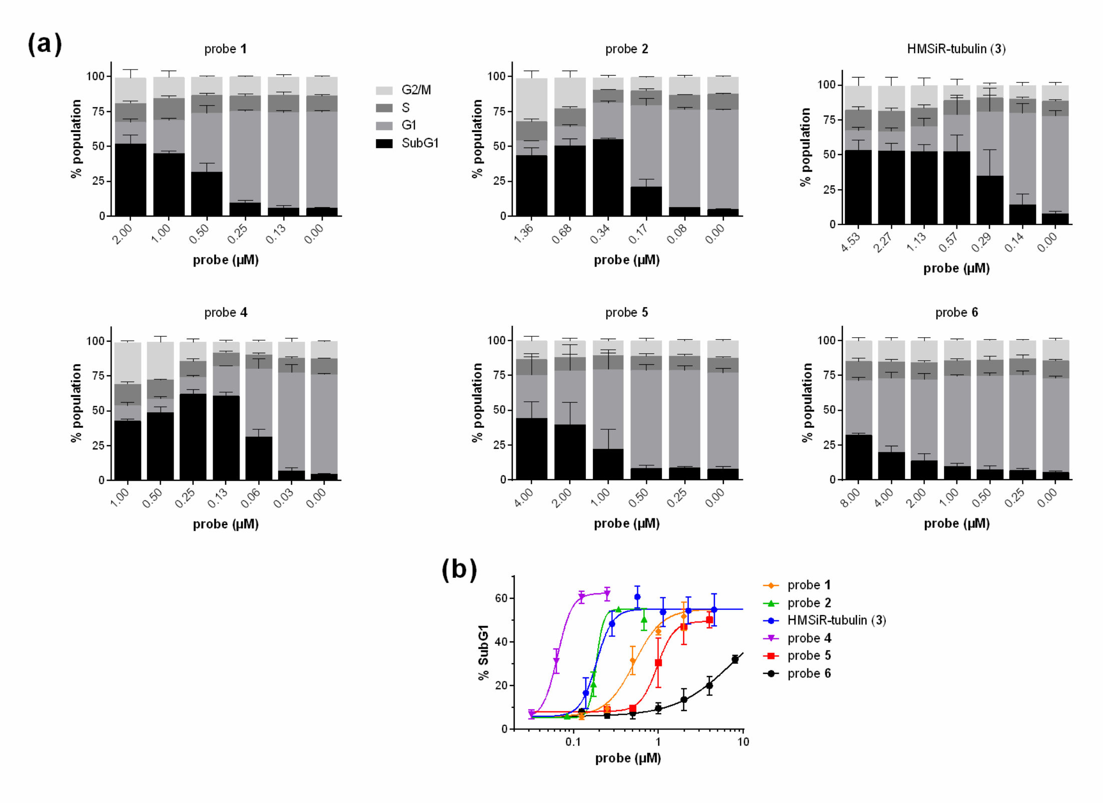

**Figure S2. Effect of the HMSiR based tubulin probes on the cell cycle of Hela cells.**

Hela cells were cultured in the presence of the indicated probes for 24h and processed according to NucleoCounter® NC-3000™ (*Chemometec*) two-step cell cycle analysis protocol as described[1]. In each sample ~10 000 cells were analyzed and the obtained cell cycle histograms were processed with ChemoMetec NucleoView NC-3000 software, version 2.1.25.8. **(a)** Stacked bar diagrams representing distribution of population among different cell cycle stages. All experiments were repeated 3-4 times and the results are presented as mean ± SD. **(b)** Data fitting into four parameter dose response equation (see Methods section for details). The derived $IC_{50}$ values are presented in **Figure 2c** of the main text.

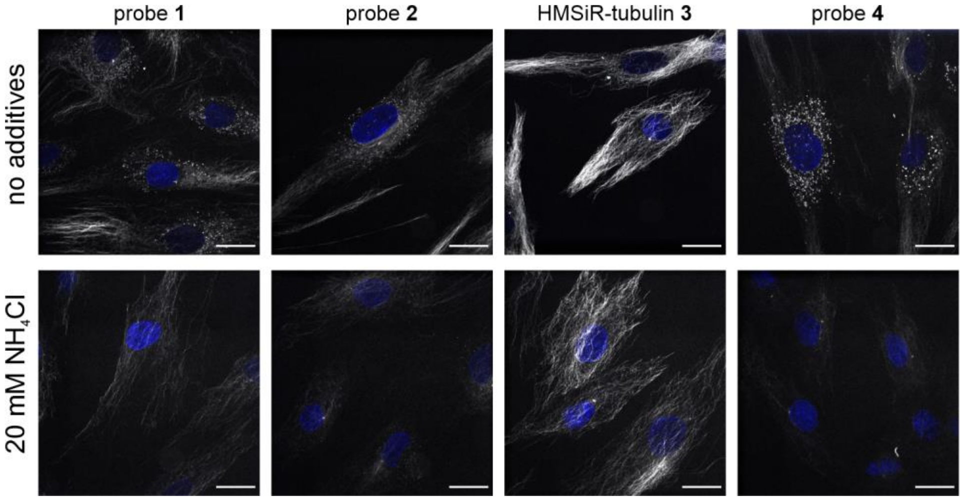

**Figure S3. Increase in lysosomal pH prevents vesicle staining by HMSiR-CTX probes.**

Living human fibroblasts were stained with 300 nM probe and 0.1 µg/ml Hoechst 33342 in DMEM medium with FBS for 1h at 37°C, then $NH_4Cl$ was added to 20 mM concentration and the cells were imaged on a spinning disk confocal microscope. Maximal intensity projections of 39 planes (living cells) acquired with a step size of 200 nm, are shown. Probe is represented in grayscale, Hoechst 33342 is blue. Scale bar – 20 µm.

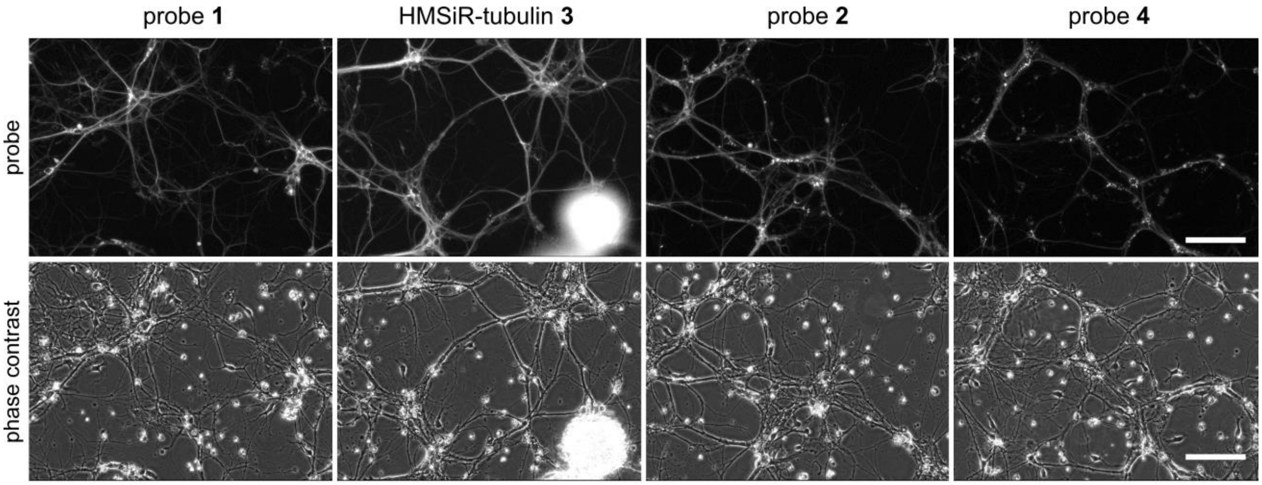

**Figure S4. Staining of mouse primary hippocampal neurons with HMSiR-CTX probes.**

The neurons were stained in the growth medium with 300 nM probe for 1h at 37°C and imaged without washing on a wide-field Lionheart FX Automated Microscope (Biotek) with 40× objective in Cy5 channel (excitation 623 nm, filter cube 628/685 nm). Scale bar – 100 µm.

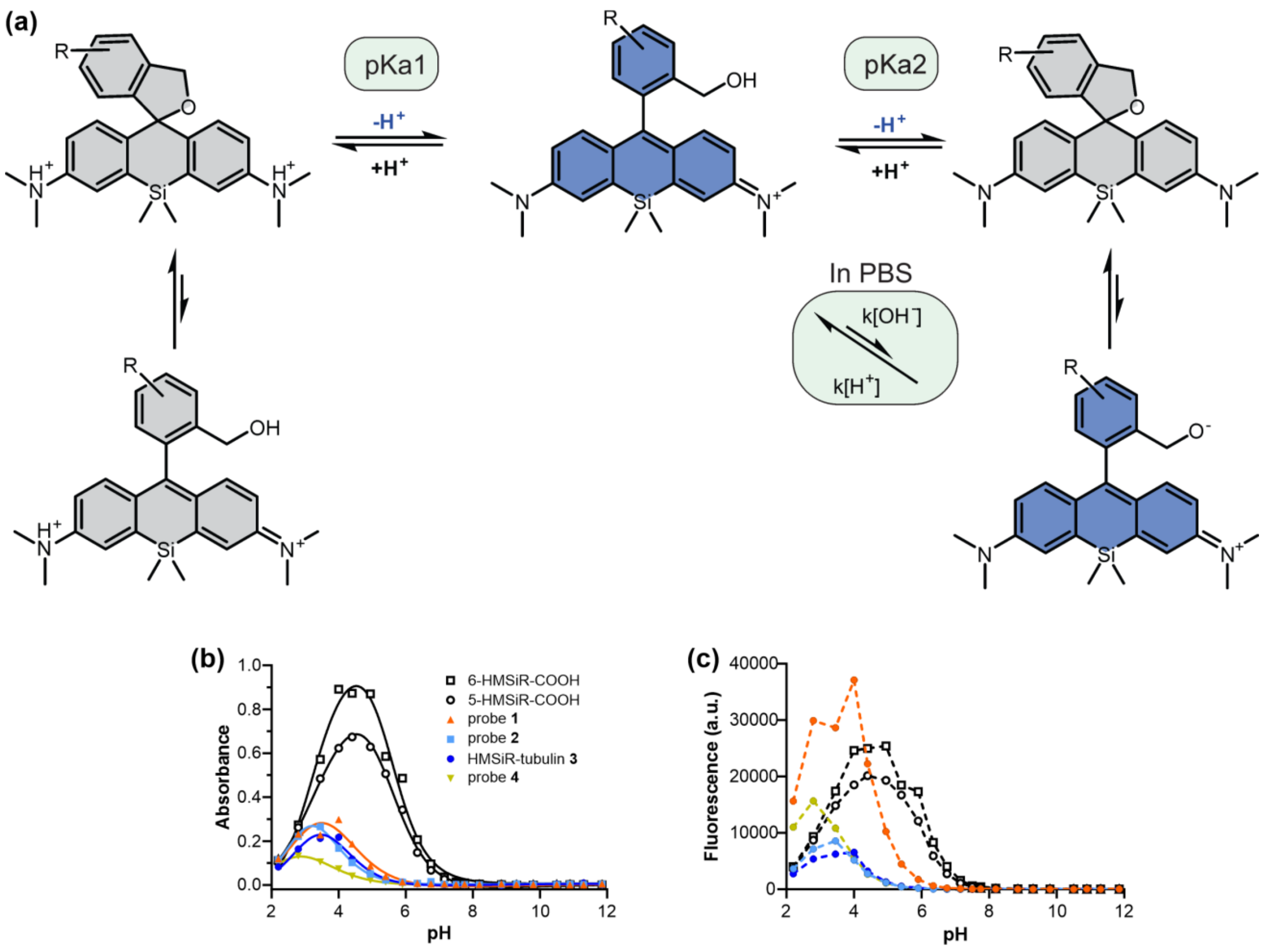

**Figure S5. Effect of pH on absorbance and fluorescence of HMSiR-CTX probes.**

**(a)** Graphical representations of pH-dependent transitions between different plausible states of HMSiR dye with depicted fluorescent-colored (blue) and non-fluorescent-colorless (gray) states. Absorbance **(b)** and fluorescence **(c)** of as a function of pH. $pK_{cyc}^{app}$ values derived from fitting absorbance data is presented in **Figure 2e**. Note that in case the fluorescence data, the fitting did not converge, likely due to a complex effect of aggregation on fluorescence.

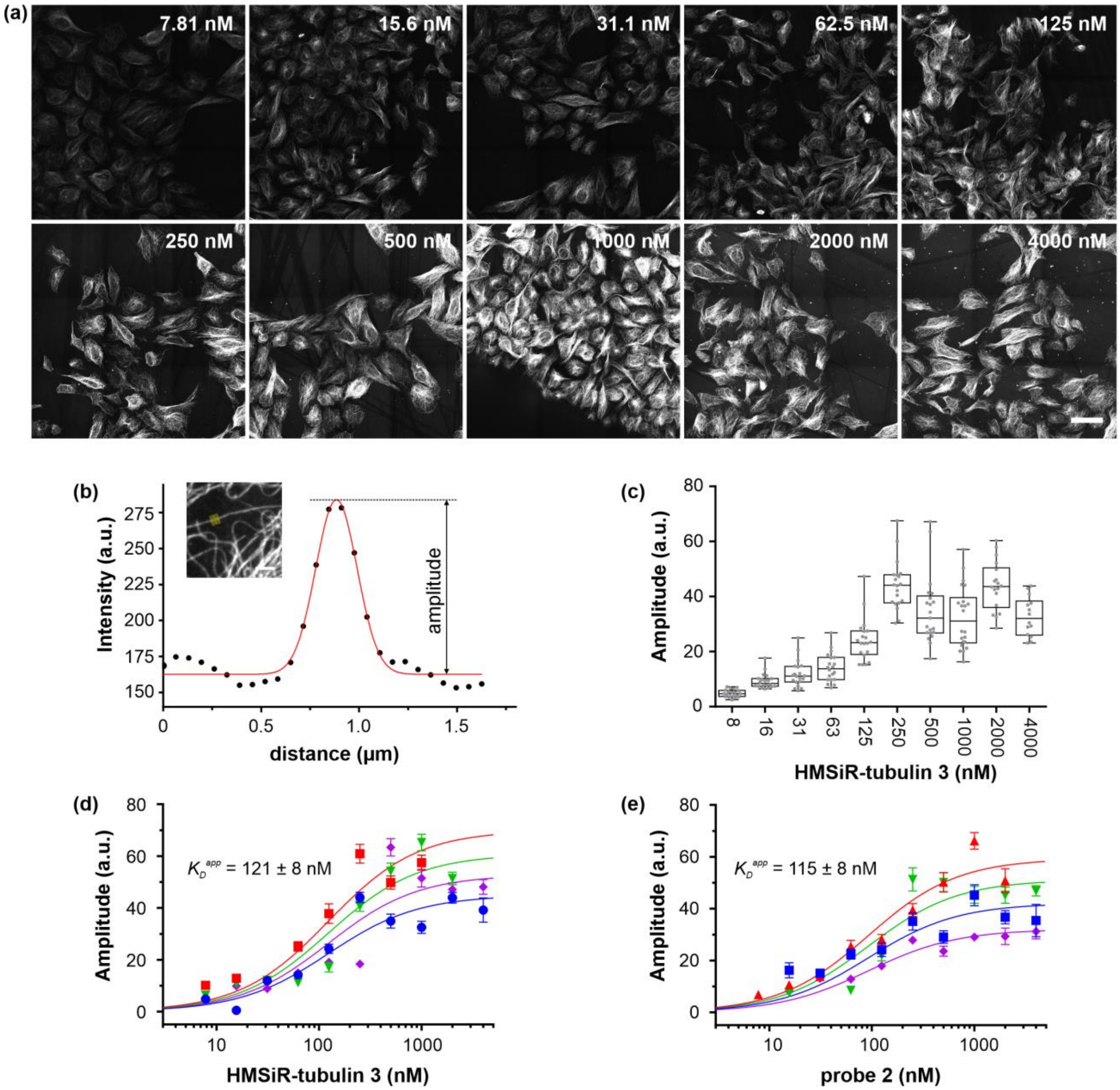

**Figure S6. Determination of probe affinity to a single microtubule on fixed U-2 OS cells.**

**(a)** Cell staining with a serial 2-fold dilutions of HMSiR-tubulin **3**. Maximum intensity projections of 41 200 nm z-slices are shown. Scale bar – 50 µm. **(b)** Line profile over 1 µm straight region of a microtubule was drawn, the resulting profile was fitted to an amplitude-version of Gaussian function, and the derived amplitudes were plotted against probe concentration. **(c)** An example of a single experiment quantification. The measurements from 4 independent experiments were fitted to the equation for a single site binding to yield $K_D^{app}$ for HMSiR-tubulin **3** **(d)** and probe **2** **(e)**.

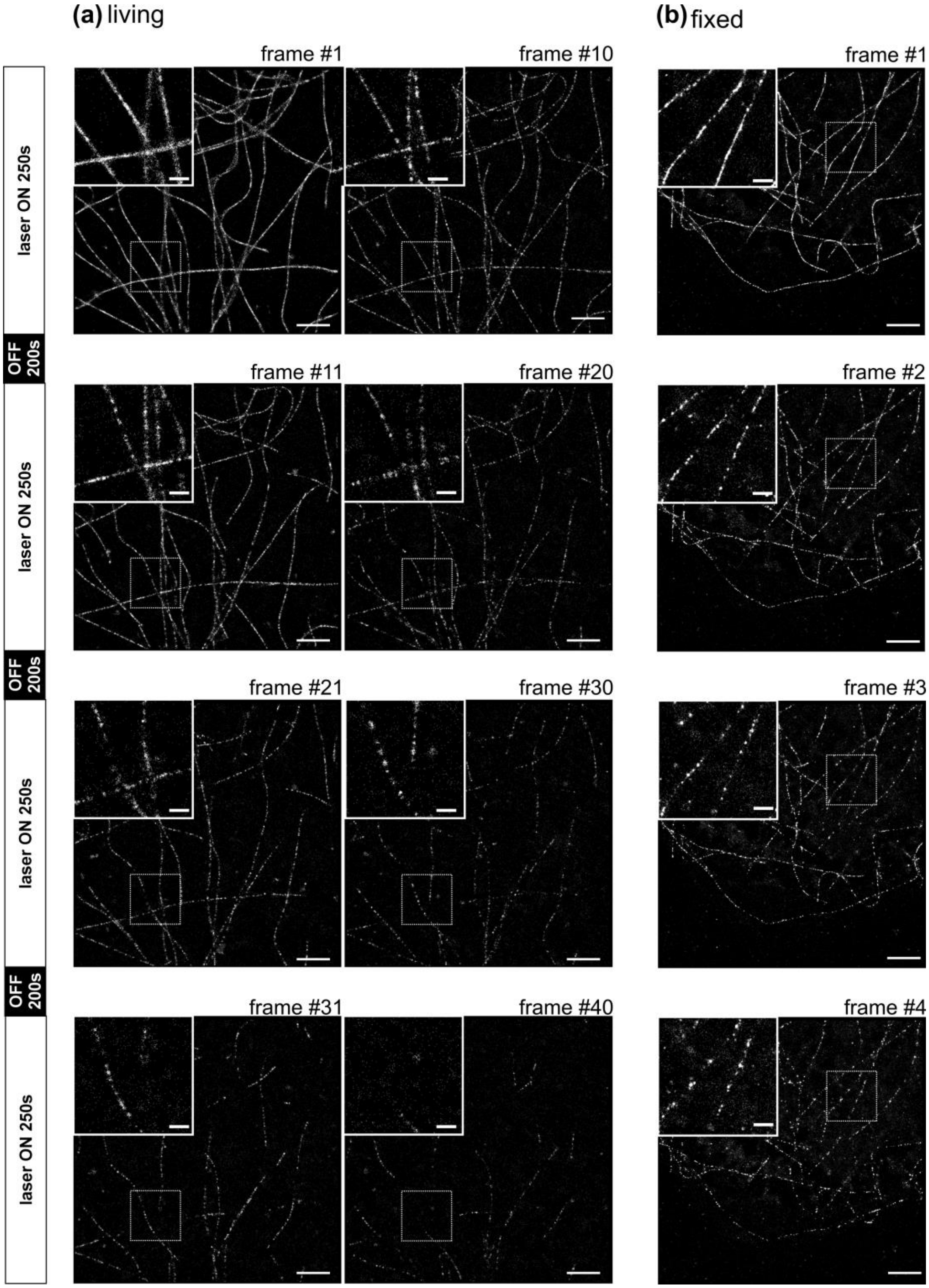

**Figure S7. Absence of fluorescence recovery after bleaching in living (a) and fixed (b) U-2 OS cells.**

The samples were illuminated with 640 nm excitation laser at 0.4 kW/cm$^2$ for 250 s, then the illumination was switched off for 200 s, and the cycle was repeated another 3 times. Living cells were imaged at 100 Hz, and a 10-frame movie was reconstructed (2500 images/frame). Note, that only the first and the last frames from each cycle are shown. Fixed cells were imaged at 20 Hz and all 4500 frames were used to reconstruct a high resolution image. Scale bar 2 µm, insets – 0.5 µm.

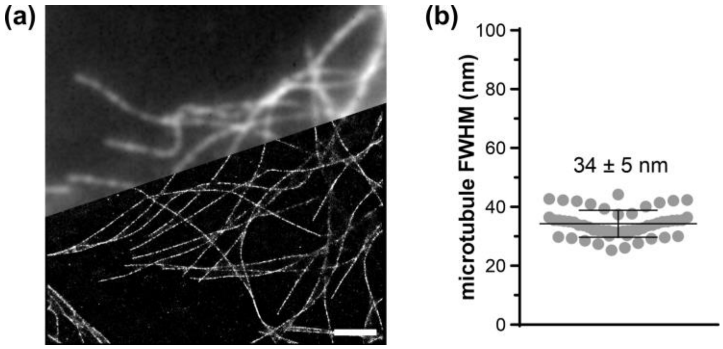

**Figure S8. SMLM imaging of tubulin in fixed U-2 OS cells stained with probe 2.**

Fixed U-2 OS cells were stained with 10 nM **2** in glycerol buffer and imaged without washing. **(a)** Comparison of diffraction-limited and SMLM image. Imaging and rendering settings are the same as for HMSiR-tubulin presented in **Figure 3b**. Scale bar 2 µm. **(b)** FWHM of microtubules, obtained from 3 fields of view.

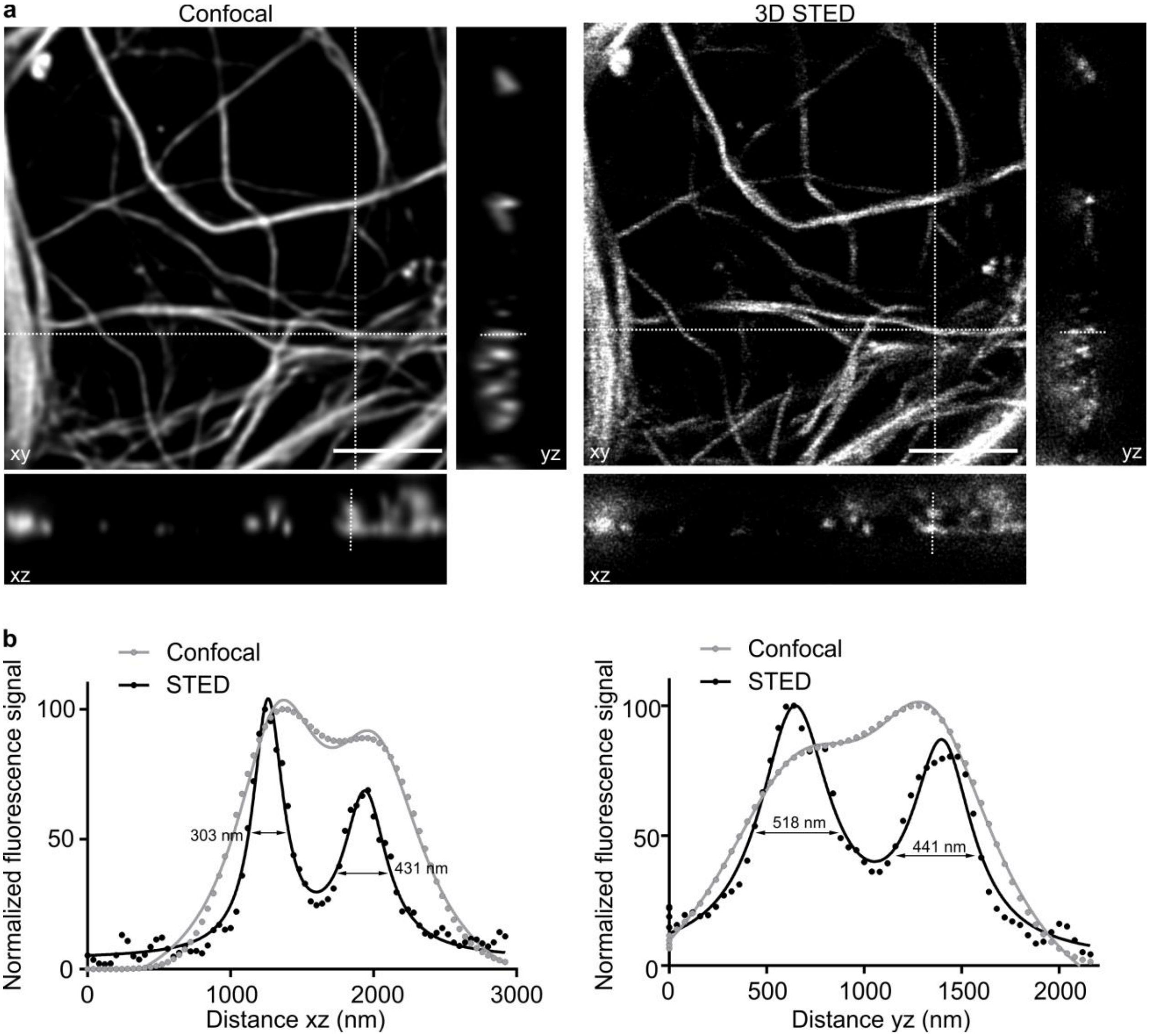

**Figure S9. 3D STED nanoscopy image of living primary mouse neurons stained with HMSiR-tubulin.**

**(a)** Comparison of confocal and 3D STED image of tubulin network stained with 300 nM probe for 1 h at 37°C. The image was acquired without probe removal and is presented as xy, xz and yz planes. Scale bar: 5 µm. **(b)** Line profile of the region marked with the dashed line in xz and yz planes of panel (a). Numbers indicate FWHM obtained by fitting data to a sum of two Lorentz distributions.

#### Supplementary movies

**Supplementary Video S1.** Time lapse series of HMSiR-tubulin (**3**) blinking in living U-2 OS cells.
The cells were stained with 100 nM probe and imaged without washing. Diffraction-limited image was generated from the movie by summing-up all frames and is shown as a reference structure in magenta.

**Supplementary Video S2.** Time lapse series of HMSiR-tubulin (**3**) blinking on fixed cytoskeleton of U-2 OS cells.
The cells were stained with 10 nM probe and imaged without washing. Diffraction-limited image was generated from the movie by summing-up all frames and is shown as a reference structure in magenta.

**Supplementary Video S3.** Microtubule dynamics in living U-2 OS cells.
The cells were stained with 100 nM HMSiR-tubulin (**3**), washed and imaged in growth medium.

**Supplementary Video S4.** Rotating maximum intensity projection of tubulin network in mouse primary neurons stained with HMSiR-tubulin (**3**).
Living cells were stained with 300 nM probe for 1h at 37°C in a growth medium and were imaged without washing. 3D STED image was acquired using 40×40×40 nm voxel size and deconvolved with SVI Huygens Essentials package.

### Supplementary materials and methods

#### Materials

Probes were dissolved in DMSO as 1000× stocks and stored at -20°C. >99% pure porcine brain tubulin (#T240) and General Tubulin buffer (#BST05) was from Cytoskeleton Inc.

#### Spectra and fluorogenicity of probes

Tubulin was dissolved in ice-cold General tubulin buffer (80 mM PIPES pH 6.9, 2 mM $MgCl_2$ and 0.5 mM EGTA) + 1 mM GTP immediately before use. Absorbance and fluorescence spectra of 10 µM probes were recorded in 100 µl volume under three conditions: i) in PBS, ii) in General Tubulin buffer + 1 mM GTP with 1 mg/ml (~20 µM) tubulin and iii) in ethanol + 0.1% trifluoroacetic acid (TFA). The samples were prepared in Corning® 96-well Half Area High Content Imaging Glass Bottom Microplates (#4580). PBS and tubulin samples were incubated for 3 h at 37°C, in order for tubulin to polymerize completely. Samples in ethanol + 0.1% TFA were incubated for 30 min. at room temperature. The spectra were acquired on Tecan Spark20M plate reader in a bottom reading mode. Excitation wavelength was 590 nm, bandwidth 15 nm; emission wavelength step size was 2 nm, emission bandwidth was 5 nm. The spectra were processed and analyzed in GraphPad Prism version 6.03 (background subtraction, spectra averaging and normalization) and Spectragryph v1.2.11 (peak determination and correction for light scattering) (F.Menges "Spectragryph - optical spectroscopy software", Version 1.2.11, 2019, http://www.effemm2.de/spectragryph/). For background correction, spectra of the solutions containing no probes, but equivalent amount of DMSO, were acquired and subtracted from the respective probe spectra. Then the absorbance and fluorescence spectra of a probe were normalized to the $A_{659\ nm}$ or $F_{673\ nm}$ of the ethanol + TFA sample.

#### Staining and imaging of living cells

U-2 OS cells were grown in McCoy's medium. For imaging, the cells were seeded in DMEM (Dulbecco's Modified Eagle Medium) supplemented with 10 % FBS (fetal serum albumin) 24-48 h before the experiment. 10-well uncoated glass bottom plates from Greiner Bio-One (#543079) were used for spinning disk confocal microscopy; µ-Slide 8 Well Glass Bottom slides (Ibidi #80827) were used for SMLM. For living cell staining, the cells were incubated for 1 h at 37°C in the growth medium containing the indicated concentration of probe and imaged without washing at room temperature. In some experiments Hoechst 33342 was included at 1 µg/ml.

#### Fixing and staining cells

U-2 OS cells were grown as before and fixed as described[2-3]. All steps were performed at room temperature. The cells were washed four times with 200 µl of PEMP (100 mM PIPES pH 6.8, 1 mM EGTA, 2 mM $MgCl_2$ and 4% PEG 8000) and permeabilized for 90 s with 0.5% Triton X-100 in PEM (PEMP without PEG 8000). The samples were washed again 4 times with 200 µl of PEMP. The cells were fixed by incubating for 15 min. with 200 µl of 0.2% glutaraldehyde in PEM, then the solution was replaced with 200 µl of 2 mg/ml $NaBH_4$ in PEM (dissolved immediately before use) and incubation continued for another 15 min. The samples were washed 4 times with 200 µl PEM.

For determining the binding affinity, the samples were incubated for ≥30 min. at room temperature with the indicated concentration of probe in PEM and imaged on spinning disk confocal microscope without washing.

For SMLM, the final washes were performed with glycerol buffer (GB, 10mM Na-$PO_4$ pH 6.8 mM EGTA 6 mM $MgCl_2$, 3.4 M glycerol). Then 10 nM probe in GB was added, samples were incubated for ~3 h at 4°C and imaged without washing.

#### Isolation and culturing of mouse primary hippocampal neurons

All animal procedures were carried out in accordance with Directive 2010/63/EU of the European Parliament and the Council on the protection of animals used for scientific purposes as well as with the German Animal Protection Law (Tierschutzgesetz der Bundesrepublik Deutschland, TierSchG).

The protocol for hippocampal neurons isolation has been previously described[4]. Briefly, hippocampi were dissected from C57BL/6N mice of mixed gender at postnatal day P0-P1, treated with trypsin (0,25%; Gibco, cat. 15090046) for 25 minutes at 37°C and subsequently manually dissociated. Cells were plated on coverslips pre-coated with poly-L-ornithine (100 µg/ml; Sigma-Aldrich, cat. P3655) and laminin (1 µg/ml; BD Bioscience, cat. 354232) and cultivated in Neurobasal medium (Gibco, cat. 21103049) supplemented with B27 serum-free supplement (2%; Gibco, cat. 17504044), GlutaMax (1×; Gibco, cat. 35050061) and Penicillin-Streptomycin (100 units/ml and 100 µg/ml respectively; Gibco, cat. 15140122).

#### Spinning disk confocal microscopy and $K_D^{app}$ determination

Spinning disk confocal imaging was performed on Visitron Spinning disk/TIRF/SMLM system (Visitron Systems GmbH, Germany) using Nikon CFI Apochromat TIRF 100×C Oil NA 1.49 objective. Spinning disk images were acquired on a Prime BSI sCMOS camera (Teledyne Photometrics), pixel size 65 nm. 300 ms exposure and 100% of 640 nm laser power was used. 4 or 9 field of view were acquired as z-stacks with 200 nm step size. Fields of view were stitched with VisiView software (version 4.5.0.13), maximal intensity projections were made and single microtubule staining was quantified with Fiji[5] as outlined in **Figure S6**. The data were fitted into equation of single site binding with GraphPad Prism (version 6.03).

$$F = \frac{F_{max} \cdot c}{K_D + c}$$

$F$ – microtubule fluorescence intensity at probe concentration $c$, $F_{max}$ – microtubule fluorescence when all binding sites are saturated, $K_D$ – apparent binding affinity.

Because $F_{max}$ varied substantially from day to day, the data from 4 independent titrations were globally fitted using shared $K_D$, but individual $F_{max}$.

#### SMLM imaging and data processing

The SMLM data was acquired on Visitron Spinning disk/TIRF/SMLM system (Visitron Systems GmbH, Germany) using Nikon CFI Apochromat TIRF 100×C Oil NA 1.49 objective and Prime 95B sCMOS camera (Teledyne Photometrics), pixel size 108 nm. Images were acquired in HILO illumination mode, with TIRF angle set to 62°. 0.4 kW/cm$^2$ of 640 nm laser was used for excitation. Living cells were imaged at 100 Hz and 2500 frames were used to reconstruct the image. Fixed cells were imaged at 20 Hz and 4500 frames were used for reconstruction.

The data was processed with SVI Huygens Localizer (version 20.10) software and the positions of the emitters were determined by fitting each fluorescence event to the 2D Gaussian distribution. The obtained localizations were loosely filtered to include only those with uncertainty <50 nm and photon count <5000 for living or <10000 for fixed cells. All histograms contained 256 bins. The drift correction was applied and high resolution images were rendered with a pixel size of 5 nm and fixed FWHM of 15 nm. Mean localization uncertainty was calculated by fitting uncertainty histogram to log-normal distribution, using Origin 8.6.0. Mean photon number per molecule per frame was calculated as the mean of all localizations in Thunderstorm[6]. Diffraction-limited images were generated by summing all the frames used for reconstruction in Fiji[5].

Tiff images were analyzed with Fiji[5]. Cross section of microtubule was marked with 3 pixels straight line. Fitting of obtained intensity profiles to Gaussian distribution and FWHM values calculation was performed with program written in Python. At least 30 events from at least 3 different images were analyzed.

#### Determination of $pK_{cyc}^{app}$

Universal buffer for UV spectrophotometry was prepared by taking 50 mL solution consisting of 0.1 M citric acid (21.01 g/l), 0.1M KH2PO4 (13.61 g/l), 0.1 M sodium tetraborate (19.07 g/l), 0.1 M Tris (12.11 g/l), 0.1 M KCl (7.46 g/l) and adjusting the pH to the required values by adding x mL of 0.4 M HCl or 0.4 M NaOH, followed by the dilution to 200 ml. Buffers in the range of pH 2-12 were prepared in 0.5 increments. 1 µl of 2mM stock solution in DMSO of the probe was distributed along the wells of 96-well plate and 300 µl of buffer at certain pH value was added, different buffers with pH increments of 0.5 were used in other wells along the pH range from 2 to 12. The plate was kept at room temperature before the measurements. Absorption was measured on a multiwell plate reader Spark® 20M (Tecan) in glass bottom 96-well plates at room temperature (25°C). Absorption of solutions was recorded from 320 nm to 850 nm with wavelength step size of 1 nm. The background absorption of the glass bottom plate was measured in wells containing only buffer with similar amount of DMSO and subtracted from the spectra of the samples. Fluorescence emission recorded from 620 nm to 850 nm (595 nm excitation) with 5 nm emission bandwidth and 2 nm step. The obtained maximum absorption or emission values were plotted against corresponding pH values in GraphPad PRISM software. The obtained graphs were fitted to a bell-shaped dose response curve, characterized by two $pK_a$ values. The larger $pK_a$ value was adopted as $pK_{cyc}^{app}$.

#### General chemical experimental information and synthesis

NMR spectra were recorded at 25 °C with an Agilent 400-MR spectrometer at 400.06 MHz (1H) and 100.60 MHz ($^{13}$C) and Bruker Avance III HD 500 spectrometer (av500) at 500.25 MHz ($^{1}$H) and 125.80 MHz ($^{13}$C) and are reported in ppm. All $^{1}$H spectra are referenced to tetramethylsilane (δ = 0 ppm) using the residual signals of the solvents according to the values reported in literature[7]. Multiplicities of signals are described as follows: s = singlet, d = doublet, t = triplet, q = quartet, p = pentet, m = multiplet or overlap of non-equivalent resonances; br = broad signal. Coupling constants (J) are given in Hz.

ESI-MS were recorded on a Varian 500-MS spectrometer (Agilent). ESI-HRMS were recorded on a MICROTOF spectrometer (Bruker) equipped with ESI ion source (Apollo) and direct injector with LC autosampler Agilent RR 1200. Liquid chromatography:

Analytical LC–MS analysis was performed on an Agilent 1260 Infinity II LC/MS system equipped with an autosampler, diode array detector WR, fluorescence detector Spectra and Infinity Lab LC/MSD 6100 series quadruple with API electrospray. Analysis was done by using an Agilent Zorbax SB-C18 RRHT, 2.1 x 50 mm, 1.8 µm threaded column and SUPELCO Titan C18, 2.1 x 75 mm, 1.9 µm column with A: 25 mM $HCOONH_4$ (pH = 3.6) aqueous buffer and B: MeOH

Preparative HPLC was performed on an Interchim puriFlash 4250 2X preparative HPLC/Flash hybrid system (Article No. 1I5140, Interchim) with a 2 mL / 5 mL injection loop, a 200-600 nm UV-Vis detector and an integrated ELSD detector (Article No. 1A3640, Interchim). Preparative column: Eurospher II 100-5 C18 5 µm, 250×20.0 mm (Article No.: 25PE181E2J, Knauer), typical flow rate: 25 mL/min, unless specified otherwise. Analytical TLC was performed on Merck Millipore ready-to-use plates with silica gel 60 (F254) (Cat. No. 1.05554.0001). Flash chromatography was performed on Biotage Isolera flash purification system using the indicated type of cartridge and solvent gradient.

Source of important chemical reagents used in the study: **Cabazitaxel** was bought from Carbosynth. **Docetaxe**l was bought from TCI. **Larotaxel** was synthesized according to the previously described procedure[8]. **6-HMSiR-COOH** and **5-HMSiR-COOH** were synthesized according to previously published procedure[9].

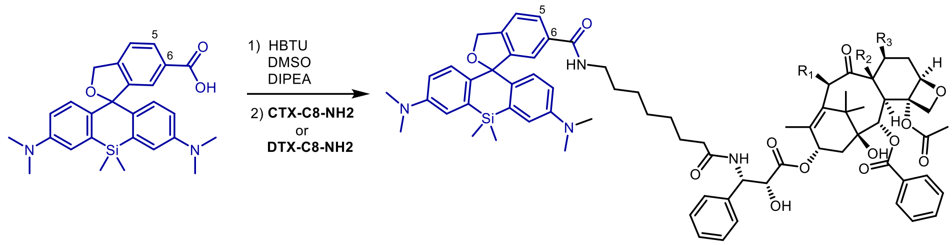

Probe **4**, $R_1$ = -$OCH_3$; $R_2$ = -$CH_3$; $R_3$ = -$OCH_3$.
Probe **5**, $R_1$ = -OH; $R_2$ = -$CH_3$: $R_3$ = -OH

**Scheme S1.** Synthetic route of probes **4** and **5.**

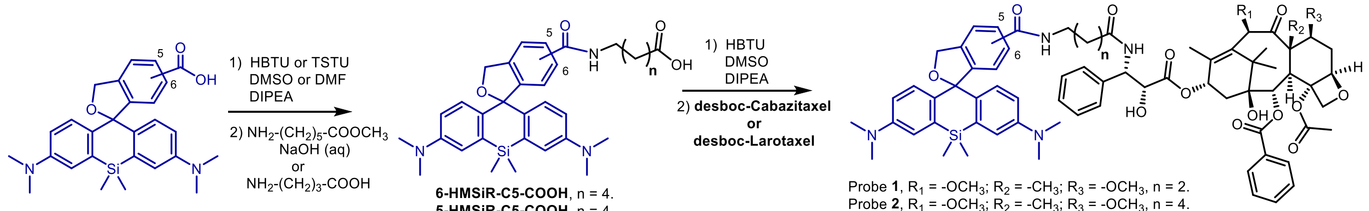

**6-HMSiR-C5-COOH**, n = 4.
**5-HMSiR-C5-COOH**, n = 4.
**6-HMSiR-C3-COOH**, n = 2.

Probe **1**, $R_1$ = -$OCH_3$; $R_2$ = -$CH_3$; $R_3$ = -$OCH_3$, n = 2.
Probe **2**, $R_1$ = -$OCH_3$; $R_2$ = -$CH_3$; $R_3$ = -$OCH_3$, n = 4.
Probe **3** (*HMSiR-tubulin*) , $R_1$ = -$OCH_3$; $R_2$ = -$CH_3$; $R_3$ = -$OCH_3$, n = 4.
Probe **6**, $R_1$ = -OAc; $R_2$ = $R_3$ = -$CH_2$-. n = 4.

**Scheme S2.** Synthetic route of probes **1-3** and **6**.

Synthesis procedure for **6-HMSiR-C5-COOH** and **5-HMSiR-C5-COOH:**

**6-HMSiR-COOH** or **5-HMSiR-C5-COOH** (0.0222 mmol, 10 mg, 1 eq.), DIPEA (38 µL, 0.222 mmol, 10 eq.) and TSTU (0.0333 mmol, 10 mg, 1.5 eq.) were dissolved in 500 µL of dry DMF and stirred at room temperature for 1 hour. A solution of methyl 6-aminohexanoate hydrochloride (0.0444 mmol, 8.1 mg, 2 eq.) in 400 µL of DMFwas added to the reaction mixture and stirring continued for 2 hours. Then 500 µL of 1M NaOH solution was added and reaction mixture was stirred overnight. Reaction mixture was quenched by addition of 250 µL of formic acid. Solvents were evaporated under reduced pressure, the leftover was redissolved in MeCN : $H_2O$ mixture and product was purified by preparative HPLC (preparative column: Eurospher 100 C18, 5 µm, 250 × 20 mm; solvent A: acetonitrile, solvent B: H2O + 0.2% v/v HCOOH; temperature 25 °C, Purified product lyophilized from acetonitrile:water or 1,4-dioxane:water mixture to obtain 7 mg (55%) of **6-HMSiR-C5-COOH** or 8.5 mg (67%) of **5-HMSiR-C5-COOH** as light blue powders.

**6-HMSiR-C5-COOH:**

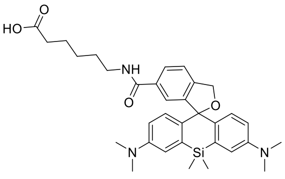

6-HMSiR-C5-COOH

$^1$H NMR (400 MHz, $d_6$-dmso) δ 11.88 (s, 1H), 8.31 (t, *J* = 5.7 Hz, 1H), 7.67 (dd, *J* = 7.9, 1.5 Hz, 1H), 7.39 (d, *J* = 7.9 Hz, 1H), 7.23 (d, *J* = 1.5 Hz, 1H), 7.01 (d, *J* = 9.0 Hz, 2H), 6.90 (d, *J* = 2.9 Hz, 2H), 6.65 (dd, *J* = 9.0, 2.9 Hz, 2H), 5.41 (s, 2H), 3.10 (q, *J* = 6.7 Hz, 2H), 2.86 (s, 12H), 2.13 (t, *J* = 7.4 Hz, 2H), 1.48 – 1.36 (m, 4H), 1.23 – 1.18 (m, 2H), 0.60 (s, 3H), 0.44 (s, 3H).

ESI-MS, positive mode: m/z = 572.3 $[M+H]^+$.

HRMS (ESI) calcd for $C_{33}H_{42}N_3O_4Si$ $[M+H]^+$ 572.2939, found 572.2931

**5-HMSiR-C5-COOH:**

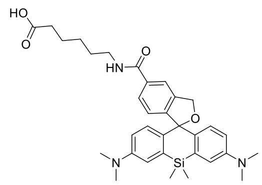

5-HMSiR-C5-COOH

$^1$H NMR (400 MHz, $CD_3OD$) δ 8.18 (s, 1H), 7.88 (d, *J* = 7.8 Hz, 1H), 7.35 (d, *J* = 2.9 Hz, 2H), 7.24 (d, *J* = 7.8 Hz, 1H), 7.03 (d, *J* = 9.6 Hz, 2H), 6.75 (dd, *J* = 9.6, 2.9 Hz, 2H), 4.34 (s, 2H), 3.44 (t, *J* = 7.1 Hz, 2H), 3.33 (s, 12H), 2.32 (t, *J* = 7.4 Hz, 2H), 1.71 – 1.64 (m, 4H), 1.49 – 1.42 (m, 2H), 0.60 (s, 3H), 0.59 (s, 3H).

ESI-MS, positive mode: m/z = 572.3 $[M+H]^+$.

HRMS (ESI) calcd for $C_{33}H_{42}N_3O_4Si$ $[M+H]^+$ 572.2939, found 572.2933.

Synthesis procedure for **6-HMSiR-C3-COOH**:

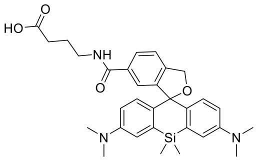

6-HMSiR-C3-COOH

**6-HMSiR-COOH** (0.0222 mmol, 10 mg, 1 eq.), DIPEA (38 µL, 0.222 mmol, 10 eq.) and HBTU (0.0266 mmol, 10 mg, 1.2 eq.) were dissolved in 500 µL of dry DMSO and stirred at room temperature for 15 min. A solution of 6-aminobutyric acid (0.0666 mmol, 6.7 mg, 3 eq.) in 500 µL of DMSO:$H_2O$ mixture (1:1) was added to the reaction mixture and stirring continued for 1 hour. Reaction mixture was quenched with 100 µL of formic acid. Solvents were evaporated under reduced pressure, the leftover was redissolved in MeCN:$H_2O$ mixture and product was purified by preparative HPLC (preparative column: Eurospher 100 C18, 5 µm, 250 × 20 mm; solvent A: acetonitrile, solvent B: H2O + 0.2% v/v HCOOH; temperature 25 °C, gradient A:B - 5 min 20:80 isocratic, 5-30 min 20:80 to 80:20 gradient). Product lyophilized from acetonitrile: water mixture to obtain 7.8 mg (65%) of light blue powder.

$^1$H NMR (400 MHz, $d_6$-dmso) δ 11.97 (s, 1H), 8.36 (t, *J* = 5.6 Hz, 1H), 7.68 (dd, *J* = 7.9, 1.5 Hz, 1H), 7.40 (d, *J* = 7.9 Hz, 1H), 7.25 (d, *J* = 1.5 Hz, 1H), 7.02 (d, *J* = 8.9 Hz, 2H), 6.91 (d, *J* = 2.8 Hz, 2H), 6.66 (dd, *J* = 8.9, 2.8 Hz, 2H), 5.41 (s, 2H), 3.14 (q, *J* = 6.6 Hz, 2H), 2.86 (s, 12H), 2.17 (t, *J* = 7.3 Hz, 2H), 1.63 (p, *J* = 7.2 Hz, 2H), 0.60 (s, 3H), 0.44 (s, 3H).

ESI-MS, positive mode: m/z = 544.3 $[M+H]^+$.

HRMS (ESI) calcd for $C_{31}H_{38}N_3O_4Si$ $[M+H]^+$ 544.2626, found 544.2628.

Synthesis procedure of **DTX-C8-NHBoc**:

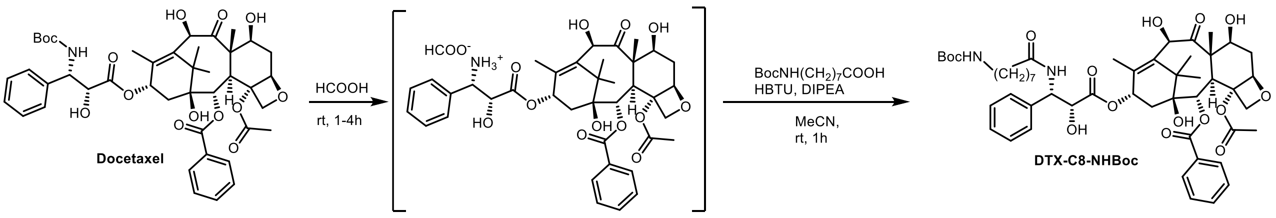

A solution of **Docetaxel** (0.124 mmol, 100 mg) in 95% formic acid (2 mL) was stirred at room temperature for 4h. Reaction progress was monitored by HPLC analysis. Once reaction was complete, formic acid was evaporated on rotary evaporator and the residue was dissolved in water and lyophilised to obtain white powder the procedure was repeated 3 times to get rid of the residual formic acid. In another flask to a solution of Boc protected 8-aminooctanoic acid (1.4 eq., 0.336 mmol, 91.7 mg) in MeCN (2 mL) was added HBTU (1.2 eq., 0.288 mmol, 109 mg), followed by addition of DIPEA (4.3 eq., 1 mmol,

100µL). Mixture was stirred at room temperature for 5 min and previously obtained **N-desboc-Docetaxel** (0.124 mmol) was added. Mixture was stirred for 1 hour. Then solvent was removed by rotary evaporator. The residue was redissolved in 70% MeCN-$H_2O$ mixture, microfiltered through a 0.45 µm PTFE membrane filter and purified by the preperative HPLC (preparative column: Knauer 100 C18, 5 µm, 250 × 30 mm; solvent A: acetonitrile, solvent B: H2O + 0.2% v/v HCOOH; temperature 25 °C, gradient A:B - 5 min 50:50 isocratic, 5-30 min 50:50 to 100:0 gradient). Fractions containing the product were collected and solvent was removed, obtained residue was lyophilised from 1:1 MeCN:$H_2O$ mixture to obtain 61 mg (52%) of

**DTX-C8-NHBoc** as a white solid.

$^1$H NMR (400 MHz,$d_6$-dmso) δ 8.36 (d, J = 9.0 Hz, 1H), 7.99 (dt, J = 7.0, 1.4 Hz, 2H), 7.69 (tt, J = 7.4, 1.4 Hz, 1H), 7.60 (t, J = 7.5 Hz, 2H), 7.39 – 7.29 (m, 4H), 7.21 (tt, J = 7.2, 1.5 Hz, 1H), 6.73 (t, J = 5.7 Hz, 1H), 5.95 – 5.88 (m, 2H), 5.42 (d, J = 7.2 Hz, 1H), 5.27 (dd, J = 9.0, 5.9 Hz, 1H), 5.10 (s, 1H), 5.01 (d, J = 7.2 Hz, 1H), 4.97 – 4.88 (m, 2H), 4.58 (s, 1H), 4.41 (t, J = 6.3 Hz, 1H), 4.09 – 3.99 (m, 3H), 3.69 (d, J = 7.2 Hz, 1H), 2.87 (q, J = 6.7 Hz, 2H), 2.28 (s, 1H), 2.24 (s, 3H), 2.15 (t, J = 7.3 Hz, 2H), 1.98 (dd, J = 15.3, 9.2 Hz, 1H), 1.82 (dd, J = 15.4, 8.9 Hz, 1H), 1.75 (s, 3H), 1.73 – 1.59 (m, 1H), 1.53 (s, 3H), 1.49 – 1.43 (m, 2H), 1.36 (s, 9H), 1.34 – 1.26 (m, 2H), 1.18 (q, J = 9.0, 7.1 Hz, 6H), 1.03 (s, 3H), 0.99 (s, 3H).

$^{13}$C NMR (101 MHz, $d_6$-dmso) δ 209.3, 172.6, 171.9, 169.7, 165.3, 155.5, 139.7, 136.8, 135.9, 133.3, 130.0, 129.6, 128.7, 128.1, 127.14, 127.10, 83.7, 80.3, 77.3, 76.9, 75.5, 74.8, 73.8, 73.6, 70.8, 70.0, 57.0, 55.0, 46.0, 42.9, 39.8, 36.5, 35.4, 35.1, 29.5, 28.6, 28.5, 28.3, 26.5, 26.2, 25.4, 22.4, 20.8, 13.7, 9.8.

ESI-MS, positive mode: m/z = 949.5 $[M+H]^+$.

HRMS (ESI) calcd for $C_{51}H_{69}N_2O_{15}$ $[M+H]^+$ 949.4692, found 949.4683.

Synthesis procedure for probes **1-3** and **6**:

A solution of **Cabazitaxel** or **Larotaxel**[8] (0.015 mmol) in 95% formic acid (1 mL) was stirred at room temperature for 1-3h. Reaction progress was monitored by HPLC analysis. Once reaction was complete, formic acid was evaporated on a rotary evaporator and residue was dissolved in water and lyophilized to obtain white powder. The lyophilisation was repeated 2-3 times in order to remove residual formic acid. The obtained compounds **N-*desboc*-Cabazitaxel** or **N-*desboc*-Larotaxel** were used further without any additional purifications.

In another flask **6-HMSiR-C5-COOH** or **5-HMSiR-C5-COOH** or **6-HMSiR-C3-COOH** (0.01 mmol, 1eq), DIPEA (52 µL, 0.3 mmol, 30 eq.) and HBTU (0.012 mmol, 4.5 mg, 1.2 eq.) were dissolved in 400 µL of dry DMSO and stirred at room temperature for 15 min. Then a solution of previously obtained **N-*desboc*-**

**Cabazitaxel** or **N-*desboc*-Larotaxel** (0.015 mmol, 1.5 eq) in 100-200 µL DMSO and 10 µL of DIPEA were added to the reaction mixture. Stirring was continued for 1 hour. Reaction progress was monitored by HPLC analysis. Reaction mixture was quenched by addition of 100 µL of formic acid. The solvents were evaporated at reduced pressure. The leftover was redissolved in MeCN:$H_2O$ mixture and purified by preparative HPLC (preparative column: Eurospher 100 C18, 5 µm, 250 × 20 mm; solvent A: acetonitrile, solvent B: $H_2O$ + 0.2% v/v HCOOH; temperature 25 °C; gradient A:B - 5 min 30:70 isocratic, 5-30 min 30:70 to 100:0 gradient). Fractions containing the products were collected, solvent evaporated under reduced pressure and leftover lyophilized from MeCN:$H_2O$ mixture.

Probe **1**:

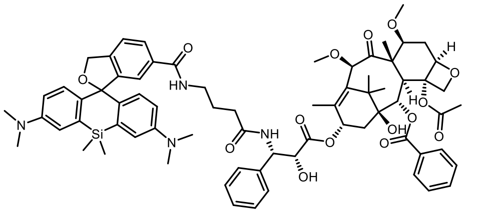

5.2 mg (41% yield) of light blue solid obtained after lyophilisation.

$^1$H NMR (400 MHz, $d_6$-dmso) δ 8.41 (d, *J* = 9.0 Hz, 1H), 8.35 (t, *J* = 5.6 Hz, 1H), 7.97 – 7.91 (m, 2H), 7.67 (dd, *J* = 7.9, 1.5 Hz, 1H), 7.63 (tt, *J* = 7.4, 2.3 Hz, 1H), 7.56 (t, *J* = 7.4 Hz, 2H), 7.39 (d, *J* = 8.0 Hz, 1H), 7.34 – 7.25 (m, 5H), 7.18 (tt, *J* = 7.1, 1.3 Hz, 1H), 7.01 (dd, *J* = 8.9, 1.5 Hz, 2H), 6.90 (d, *J* = 2.9 Hz, 2H), 6.64 (dd, *J* = 8.8, 2.8 Hz, 2H), 5.99 (d, *J* = 6.6 Hz, 1H), 5.90 (t, *J* = 9.0 Hz, 1H), 5.40 (s, 2H), 5.35 (d, *J* = 7.2 Hz, 1H), 5.22 (dd, *J* = 9.0, 6.0 Hz, 1H), 4.92 (dd, *J* = 9.6, 1.6 Hz, 1H), 4.66 (s, 1H), 4.60 (s, 1H), 4.36 (t, *J* = 5.5 Hz, 1H), 3.99 (s, 2H), 3.72 (dd, *J* = 10.6, 6.6 Hz, 1H), 3.59 (d, *J* = 7.1 Hz, 1H), 3.26 (s, 3H), 3.17 (s, 3H), 3.11 (q, *J* = 7.3 Hz, 2H), 2.86 (s, 12H), 2.67 – 2.56 (m, 1H), 2.19 (s, 3H), 2.14 (t, *J* = 7.5 Hz, 2H), 2.01 – 1.94 (m, 1H), 1.84 – 1.75 (m, 4H), 1.63 (p, *J* = 6.9 Hz, 2H), 1.49 (s, 4H), 0.96 (s, 3H), 0.92 (s, 3H), 0.60 (s, 3H), 0.44 (s, 3H).

ESI-MS, positive mode: m/z = 1261.6 $[M+H]^+$.

HRMS (ESI) calcd for $C_{71}H_{85}N_4O_{15}Si$ $[M+H]^+$1261.5775, found 1261.5777.

Probe **2**:

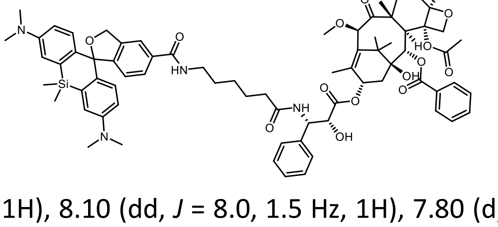

5.5 mg (42% yield) of light blue solid obtained after lyophilisation.

$^1$H NMR (500 MHz, $d_5$-pyridine) δ 9.26 (d, *J* = 9.1 Hz, 1H), 8.80 (t, *J* = 5.7 Hz, 1H), 8.37 – 8.31 (m, 2H), 8.18 (d, *J* = 0.9 Hz, 1H), 8.10 (dd, *J* = 8.0, 1.5 Hz, 1H), 7.80 (d, *J* = 7.3 Hz, 2H), 7.48 (tt, *J* = 7.1, 2.3 Hz, 1H), 7.42 (t, *J* = 7.4 Hz, 2H), 7.39 – 7.34 (m, 4H), 7.28 – 7.22 (m, 2H), 7.18 (d, *J* = 2.9 Hz, 2H), 6.85 – 6.79 (m, 1H), 6.74 (dd, *J* = 8.9, 2.9 Hz, 2H), 6.65 (s, 1H), 6.29 (dd, *J* = 9.2, 3.9 Hz, 1H), 6.24 (d, *J* = 7.1 Hz, 1H), 5.47 (s, 2H), 5.29 (d, *J* = 3.9

Hz, 1H), 5.22 (d, *J* = 8.6 Hz, 1H), 5.14 (s, 1H), 4.51 (dd, *J* = 52.8, 8.2 Hz, 2H), 4.25 (d, *J* = 7.1 Hz, 1H), 4.20 (dd, *J* = 10.6, 6.5 Hz, 1H), 3.55 (q, *J* = 6.7 Hz, 2H), 3.51 (s, 3H), 3.21 (s, 3H), 2.88 (d, *J* = 9.1 Hz, 2H), 2.81 (s, 12H), 2.80 – 2.75 (m, 1H), 2.60 (s, 3H), 2.41 (t, *J* = 6.7 Hz, 2H), 2.26 (d, *J* = 1.4 Hz, 3H), 2.11 (s, 3H), 1.92 – 1.86 (m, 1H), 1.75 (p, *J* = 7.4 Hz, 2H), 1.67 (s, 3H), 1.64 – 1.58 (m, 2H), 1.51 (s, 3H), 1.39 (p, *J* = 7.7 Hz, 2H), 0.70 (s, 3H), 0.64 (s, 3H).

ESI-MS, positive mode: m/z = 1289.6 $[M+H]^+$.

HRMS (ESI) calcd for $C_{73}H_{89}N_4O_{15}$ $[M+H]^+$ 1289.6088, found 1289.6095.

Probe **3** (*HMSiR-tubulin*)

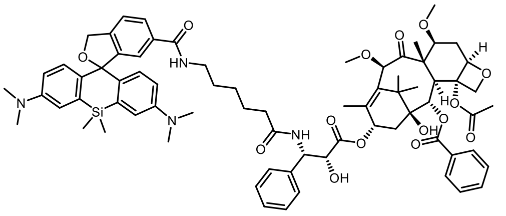

6.7 mg (52% yield) of light blue solid obtained after lyophilisation.

$^1$H NMR (500 MHz, $d_5$-pyridine) δ 9.16 (d, *J* = 9.1 Hz, 1H), 8.86 (t, *J* = 5.7 Hz, 1H), 8.32 – 8.29 (m, 2H), 8.13 (dd, *J* = 7.9, 1.5 Hz, 1H), 8.07 (d, *J* = 0.9 Hz, 1H), 7.76 (d, *J* = 7.3 Hz, 2H), 7.44 (tt, *J* = 7.3, 1.3 Hz, 1H), 7.40 – 7.37 (m, 4H), 7.36 – 7.32 (m, 2H), 7.23 (tt, *J* = 7.5, 1.1 Hz, 1H), 7.05 (t, *J* = 2.6 Hz, 2H), 6.80 – 6.73 (m, 1H), 6.63 – 6.60 (m, 2H), 6.24 (dd, *J* = 9.1, 3.9 Hz, 1H), 6.20 (d, *J* = 7.1 Hz, 1H), 5.54 (s, 2H), 5.24 (d, *J* = 3.9 Hz, 1H), 5.18 (dd, *J* = 10.0, 1.6 Hz, 1H), 5.11 (s, 1H), 4.52 (d, *J* = 8.8 Hz, 1H), 4.42 (d, *J* = 8.2 Hz, 1H), 4.21 (d, *J* = 6.8 Hz, 1H), 4.17 (dd, *J* = 10.7, 6.5 Hz, 1H), 3.56 (s, 2H), 3.48 (s, 3H), 3.38 (q, *J* = 6.8 Hz, 2H), 3.18 (s, 3H), 2.83 (d, *J* = 9.2 Hz, 2H), 2.75 (s, 6H), 2.75 (s, 6H), 2.55 (s, 3H), 2.31 – 2.25 (m, 2H), 2.21 (d, *J* = 1.4 Hz, 3H), 2.07 (s, 3H), 1.90 – 1.81 (m, 1H), 1.63 (s, 3H), 1.62 – 1.58 (m, 2H), 1.46 (s, 3H), 1.45 – 1.39 (m, 2H), 1.24 – 1.15 (m, 4H), 0.75 (s, 3H), 0.56 (s, 3H).

ESI-MS, positive mode: m/z = 1289.7 $[M+H]^+$.

HRMS (ESI) calcd for $C_{73}H_{89}N_4O_{15}Si$ $[M+H]^+$ 1289.6088, found 1289.6089.

Probe **6**

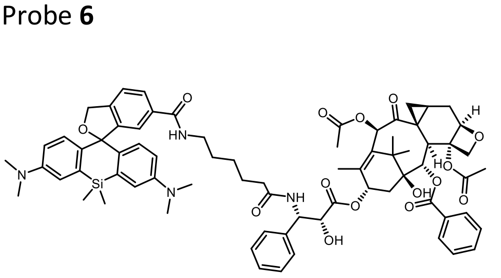

5.1 mg (40% yield) of light blue solid obtained after lyophilisation.

$^1$H NMR (400 MHz, -$d_6$-dmso) δ 8.36 – 8.27 (m, 2H), 8.01 (d, *J* = 7.2 Hz, 2H), 7.69 (dd, *J* = 8.0, 1.5 Hz, 1H), 7.65 (t, *J* = 7.2 Hz, 1H), 7.57 (t, *J* = 7.5 Hz, 2H), 7.42 (d, *J* = 8.0 Hz, 1H), 7.37 – 7.25 (m, 5H), 7.16 (t, *J* = 7.0 Hz, 1H), 7.04 (d, *J* = 8.9 Hz, 2H), 6.92 (d, *J* = 2.8 Hz, 2H), 6.67 (dd, *J* = 9.0, 2.9 Hz, 2H), 6.11 (s, 1H), 5.94 (t, *J* = 9.0 Hz, 1H), 5.86 (d, *J* = 6.7 Hz, 1H), 5.47 – 5.40 (m, 3H), 5.32 (dd, *J* = 9.2,

5.3 Hz, 1H), 4.77 (s, 1H), 4.71 (d, *J* = 2.9 Hz, 1H), 4.47 (t, *J* = 6.0 Hz, 1H), 4.01 (q, *J* = 8.5 Hz, 2H), 3.90 (d, *J* = 7.6 Hz, 1H), 3.08 (q, *J* = 6.4 Hz, 2H), 2.88 (s, 12H), 2.35 – 2.25 (m, 4H), 2.14 – 2.10 (m, 4H), 2.11 – 1.81 (m, 5H), 1.73 (s, 3H), 1.57 – 1.31 (m, 5H), 1.17 (q, *J* = 7.8, 7.3 Hz, 3H), 1.10 (s, 3H), 1.06 (s, 3H), 0.63 (s, 3H), 0.47 (s, 3H).

ESI-MS, positive mode: m/z = 1285.6 $[M+H]^+$.

HRMS (ESI) calcd for $C_{73}H_{85}N_4O_{15}Si$ $[M+H]^+$ 1285.5775, found 1285.5775.

Synthesis procedure for probe **4** and **5**:

A solution of corresponding taxane derivative **DTX-C8-NHBoc** or **CTX-C8-NHBoc**[9] (0.015 mmol) in 95% formic acid (1 mL) was stirred at room temperature for 1 h. Reaction progress was monitored by HPLC analysis. Once reaction was complete, formic acid was evaporated on a rotary evaporator and residue was dissolved in water and lyophilized to obtain white powder. The lyophilisation was repeated 2-3 times in order to remove residual formic acid. The obtained compounds were used further without any additional purifications.

The **6-HMSiR-COOH** (0.01 mmol, 4.5 mg, 1eq), DIPEA (52 µL, 0.3 mmol, 30 eq.) and HBTU (0.012 mmol, 4.5 mg, 1.2 eq.) were dissolved in 400 µL of dry DMSO and stirred at room temperature for 15 min. A solution of previously obtained deprotected taxane derivative **DTX-C8-NH$_2$** or **CTX-C8-NH$_2$** (0.015 mmol, 1.5 eq) in 100-200 µL DMSO and 10 µL of DIPEA were added to the reaction mixture. Stirring was continued for 1 hour. Reaction progress was monitored by HPLC analysis. Reaction mixture was quenched by addition of 100 µL of formic acid. The solvents were evaporated at reduced pressure. The leftover was redissolved in MeCN:$H_2O$ mixture and purified by preparative HPLC (preparative column: Eurospher 100 C18, 5 µm, 250 × 20 mm; solvent A: acetonitrile, solvent B: $H_2O$ + 0.2% v/v HCOOH; temperature 25 °C; gradient A:B - 5 min 30:70 isocratic, 5-30 min 30:70 to 100:0 gradient).) and lyophilized from MeCN:$H_2O$ mixture.

Probe **4**:

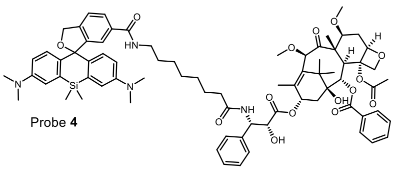

6.5mg (49% yield) of light blue solid obtained after lyophilisation.

$^1H$ NMR (400 MHz, $d_6$-dmso) δ 8.37 – 8.26 (m, 2H), 7.97 – 7.93 (m, 2H), 7.69 – 7.61 (m, 2H), 7.61 – 7.53 (m, 2H), 7.39 (d, *J* = 7.9 Hz, 1H), 7.34 – 7.26 (m, 4H), 7.24 (d, *J* = 0.8 Hz, 1H), 7.17 (tt, *J* = 7.3, 1.5 Hz, 1H), 7.01 (d, *J* = 8.9 Hz, 2H), 6.89 (d, *J* = 2.9 Hz, 2H), 6.64 (dd, *J* = 9.0, 2.9 Hz, 2H), 5.98 – 5.87 (m, 2H), 5.41 (s, 2H), 5.36 (d, *J* = 7.2 Hz, 1H), 5.25 (dd, *J* = 9.1, 5.6 Hz, 1H), 4.93 (dd, *J* = 9.6, 1.9 Hz, 1H), 4.68 (s, 1H), 4.61 (s, 1H), 4.39 (t, *J* = 5.0 Hz, 1H), 4.00 (s, 2H), 3.73 (dd, *J* = 10.6, 6.6 Hz,

1H), 3.60 (d, *J* = 7.1 Hz, 1H), 3.28 (s, 3H), 3.18 (s, 3H), 3.09 (q, *J* = 6.4 Hz, 2H), 2.86 (s, 12H), 2.69 – 2.60 (m, 1H), 2.22 (s, 3H), 2.15 – 2.07 (m, 2H), 1.99 – 1.90 (m, 1H), 1.89 – 1.81 (m, 1H), 1.80 (d, *J* = 1.4 Hz, 3H), 1.50 (s, 3H), 1.48 – 1.45 (m, 1H), 1.45 – 1.33 (m, 4H), 1.19 – 1.12 (m, 6H), 1.00 (s, 3H), 0.95 (s, 3H), 0.60 (s, 3H), 0.44 (s, 3H).

ESI-MS, positive mode: m/z = 1317.6 $[M+H]^+$.

HRMS (ESI) calcd for $C_{75}H_{93}N_4O_{15}Si$ $[M+H]^+$ 1317.6401, found 1317.6402.

Probe **5**:

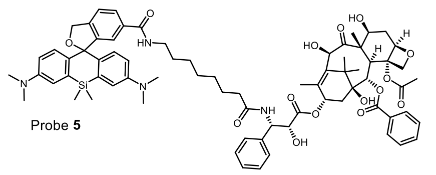
Probe **5**

5.5mg (43% yield) of light blue solid obtained after lyophilisation.

$^1$H NMR (400 MHz,$d_6$-dmso) δ 8.35 – 8.26 (m, 2H), 7.98 – 7.93 (m, 2H), 7.71 – 7.62 (m, 2H), 7.56 (t, *J* = 7.5 Hz, 2H), 7.39 (d, *J* = 8.0 Hz, 1H), 7.36 – 7.21 (m, 5H), 7.15 (t, *J* = 7.2 Hz, 1H), 7.02 (d, *J* = 8.9 Hz, 2H), 6.91 (s, 2H), 6.65 (dd, *J* = 8.9, 1.8 Hz, 2H), 5.87 (t, *J* = 8.7 Hz, 2H), 5.41 (s, 2H), 5.39 (d, *J* = 7.2 Hz, 1H), 5.23 (dd, *J* = 9.0, 5.9 Hz, 1H), 5.07 (s, 1H), 4.98 (br, s, 1H), 4.88 (dd, *J* = 9.5, 2.4 Hz, 1H), 4.55 (s, 1H), 4.37 (d, *J* = 4.5 Hz, 1H), 4.07 – 3.94 (m, 3H), 3.65 (d, *J* = 7.2 Hz, 1H), 3.54 (s, 1H), 3.09 (q, *J* = 6.8 Hz, 2H), 2.86 (s, 12H), 2.31 – 2.22 (m, 1H), 2.21 (s, 3H), 2.10 (t, *J* = 7.4 Hz, 2H), 2.00 – 1.89 (m, 1H), 1.84 – 1.73 (m, 1H), 1.72 (d, *J* = 1.4 Hz, 3H), 1.69 – 1.58 (m, 1H), 1.50 (s, 3H), 1.46 – 1.31 (m, 4H), 1.15 (d, *J* = 4.8 Hz, 6H), 0.99 (s, 3H), 0.95 (s, 3H), 0.60 (s, 3H), 0.44 (s, 3H).

ESI-MS, positive mode: m/z = 1289.6 $[M+H]^+$.

HRMS (ESI) calcd for $C_{73}H_{89}N_4O_{15}Si$ $[M+H]^+$ 1289.6088, found 1289.6087.

### NMR and HRMS spectra

#### [1]H NMR spectra copy of **6-HMSiR-C5-COOH**

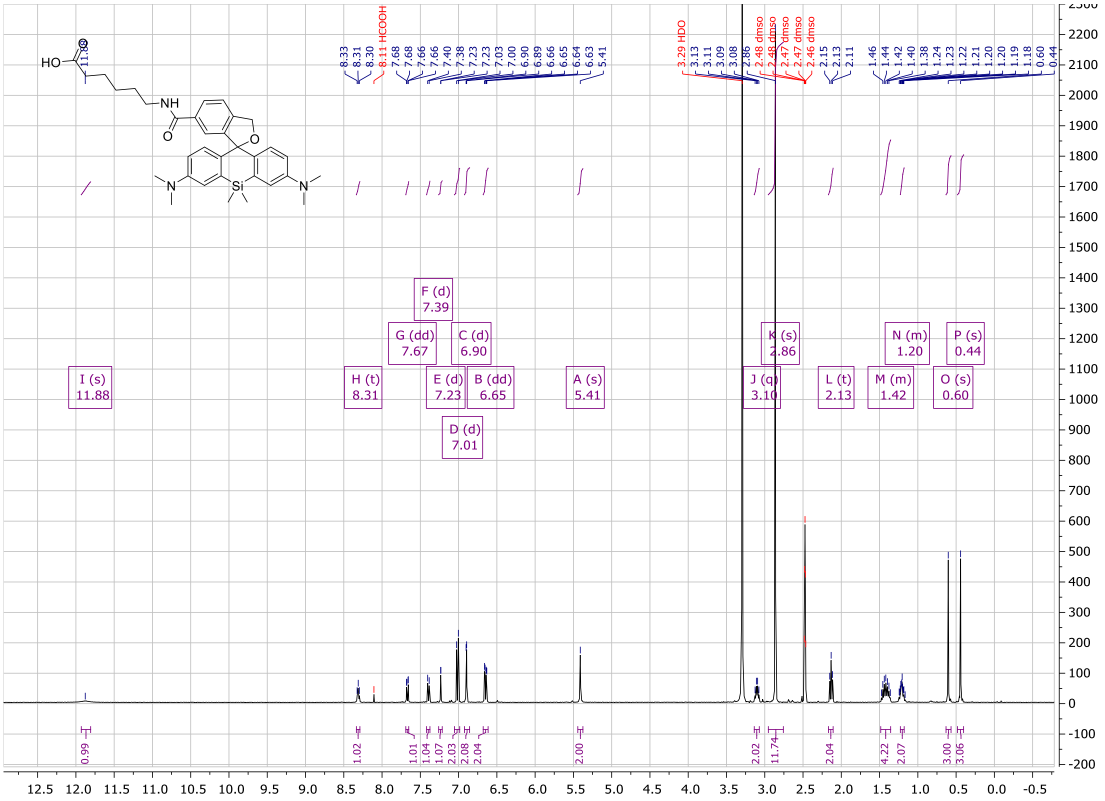

#### HRMS spectra copy of **6-HMSiR-C5-COOH**

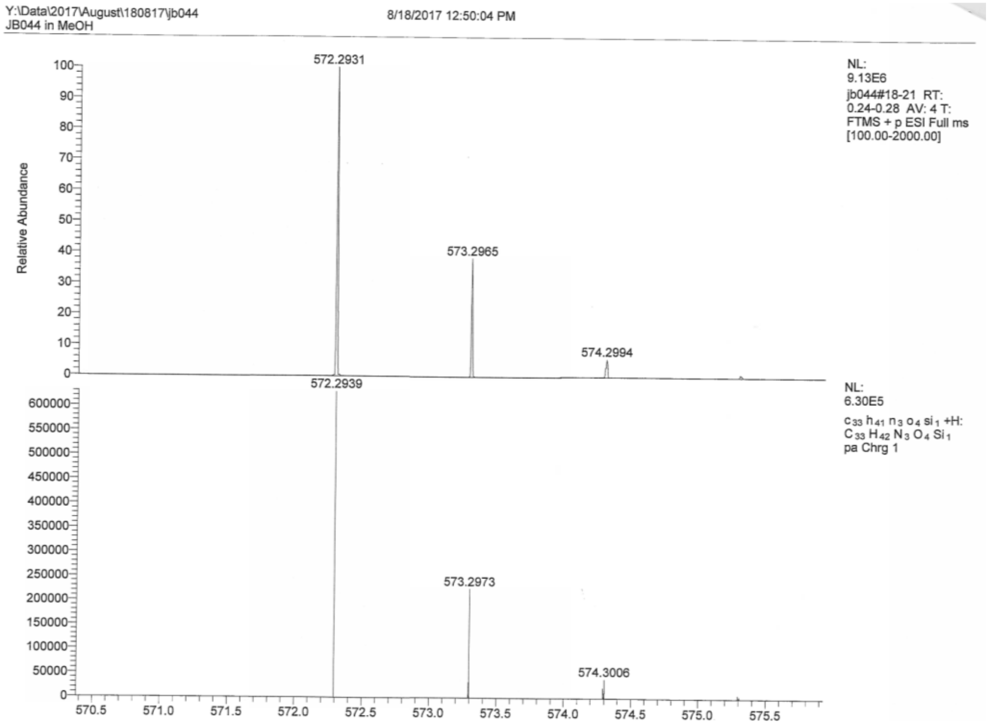

$^{1}H$ NMR spectra copy of **5-HMSiR-C5-COOH**

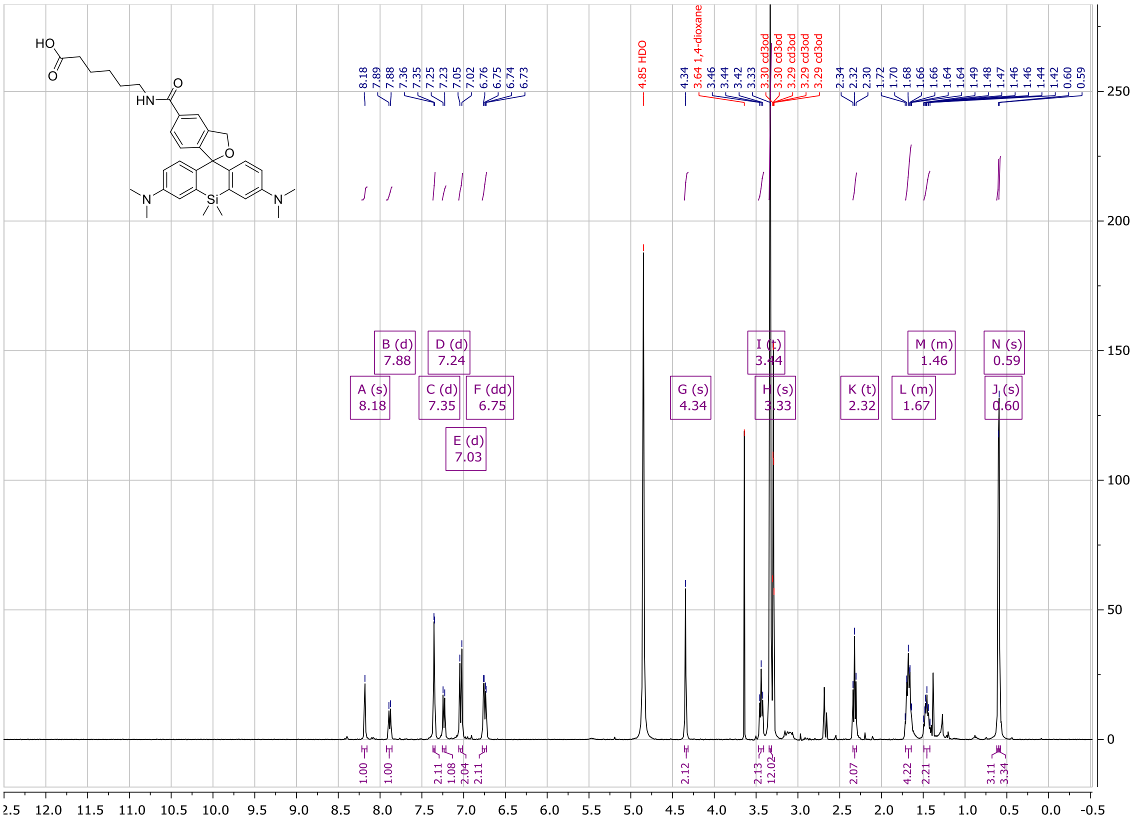

HRMS spectra copy of **5-HMSiR-C5-COOH**

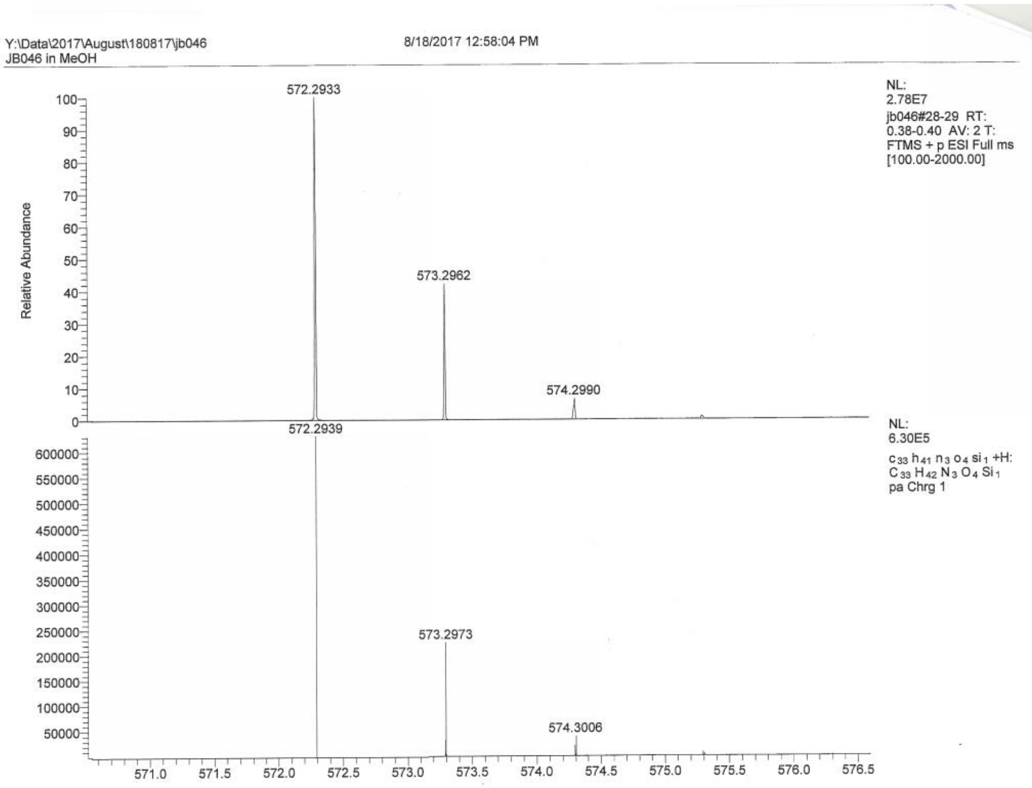

$^{1}$H NMR spectra copy of **6-HMSiR-C3-COOH**:

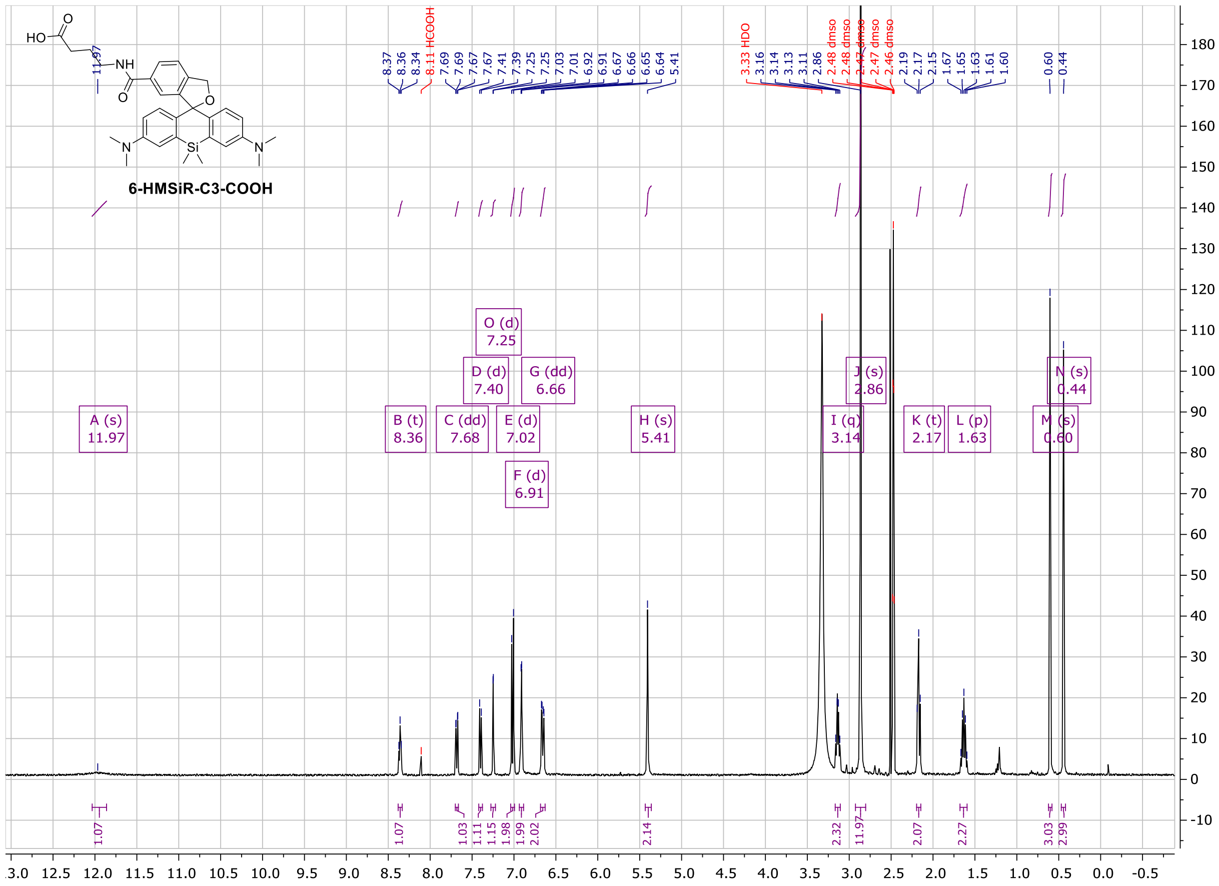

HRMS spectra copy of **6-HMSiR-C3-COOH:**

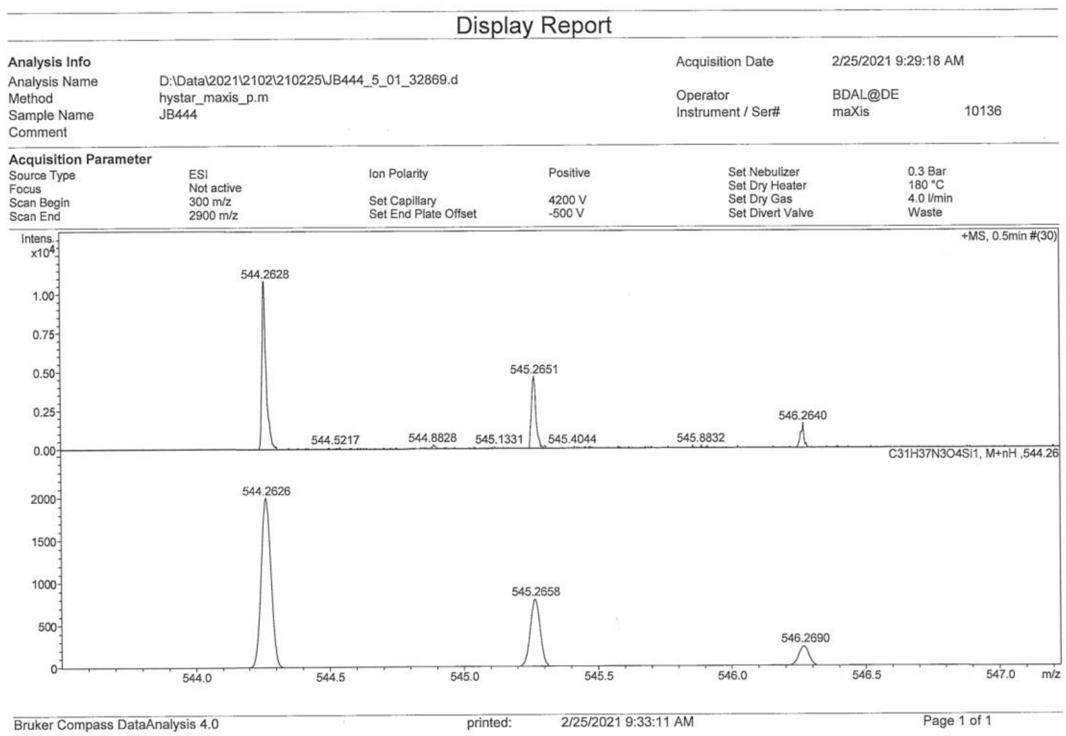

[1]H NMR spectra copy of **DTX-C8-NHBoc**

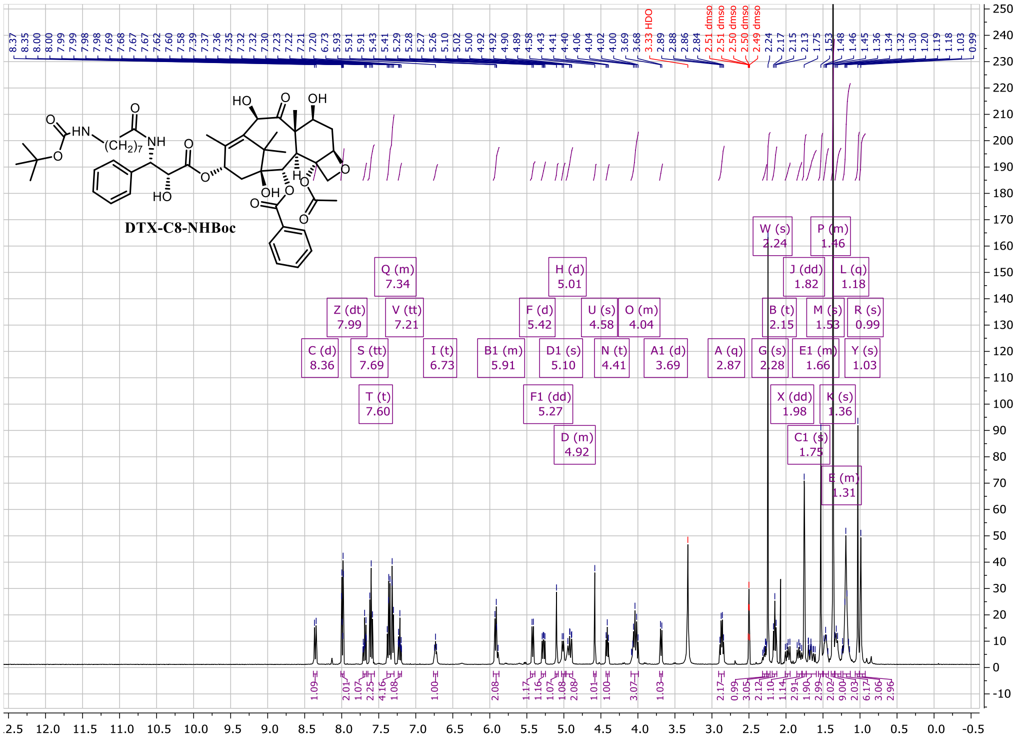

[13]C NMR spectra copy of **DTX-C8-NHBoc**

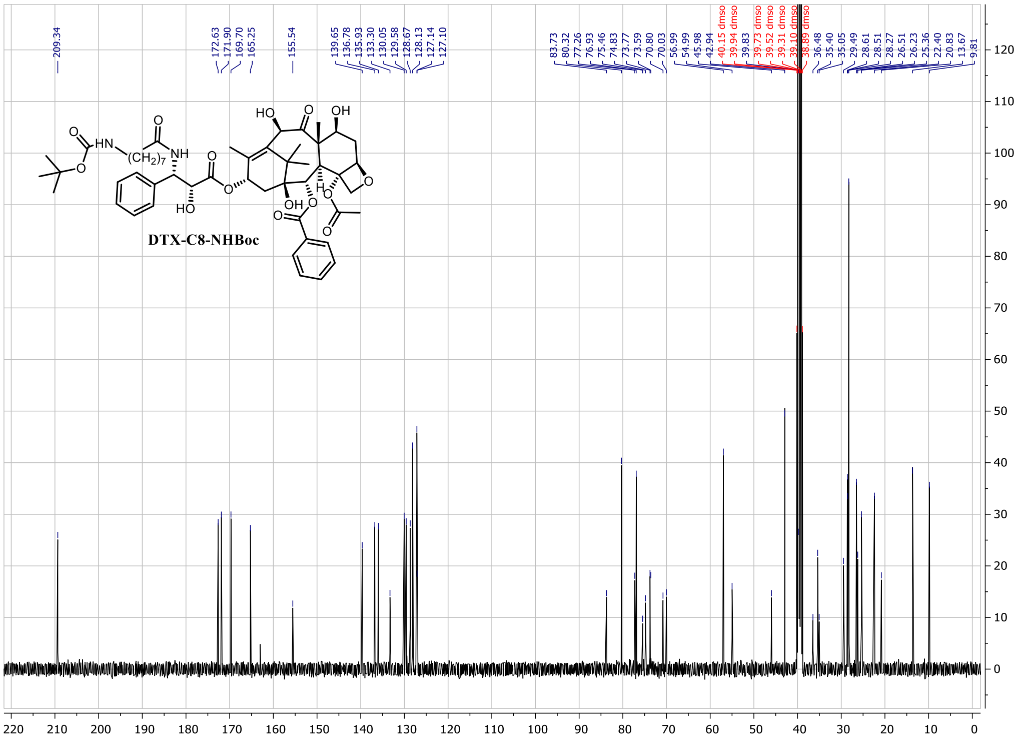

HRMS spectra copy of **DTX-C8-NHBoc**

#### Display Report

**Analysis Info**

| | | | |
|---|---|---|---|
| Analysis Name | D:\Data\2019\1909\190910\JB313_27_01_26087.d | Acquisition Date | 9/10/2019 12:14:43 PM |
| Method | hystar_maxis_p.m | Operator | BDAL@DE |
| Sample Name | JB313 | Instrument / Ser# | maXis 10136 |
| Comment | | | |

**Acquisition Parameter**

| | | | | | |
|---|---|---|---|---|---|
| Source Type | ESI | Ion Polarity | Positive | Set Nebulizer | 0.3 Bar |
| Focus | Not active | | | Set Dry Heater | 180 °C |
| Scan Begin | 300 m/z | Set Capillary | 4200 V | Set Dry Gas | 4.0 l/min |
| Scan End | 2900 m/z | Set End Plate Offset | -500 V | Set Divert Valve | Waste |

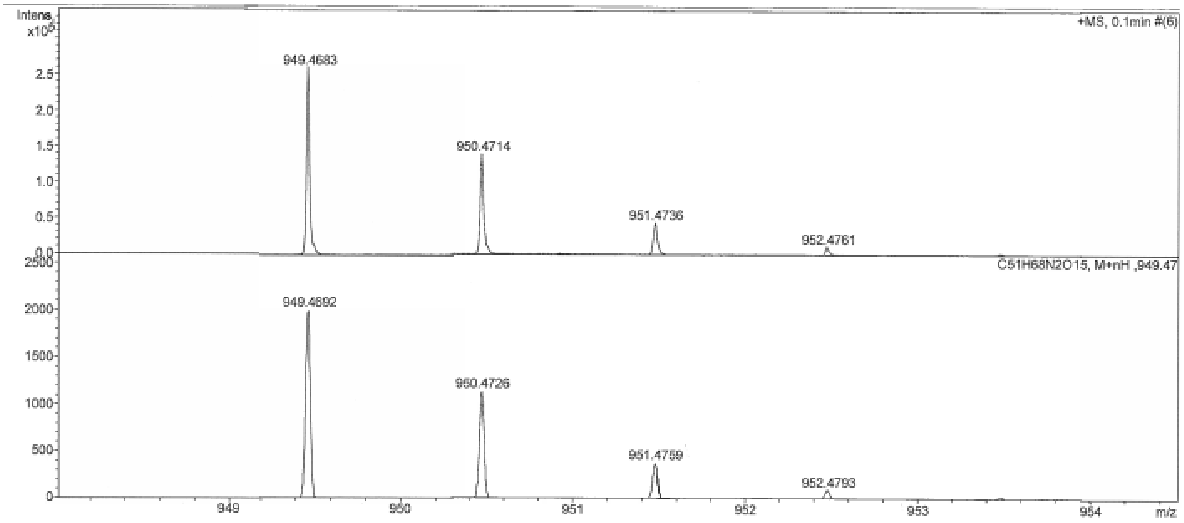

Bruker Compass DataAnalysis 4.0 printed: 9/10/2019 12:19:23 PM Page 1 of 1

[1]H NMR spectra copy of probe **5**:

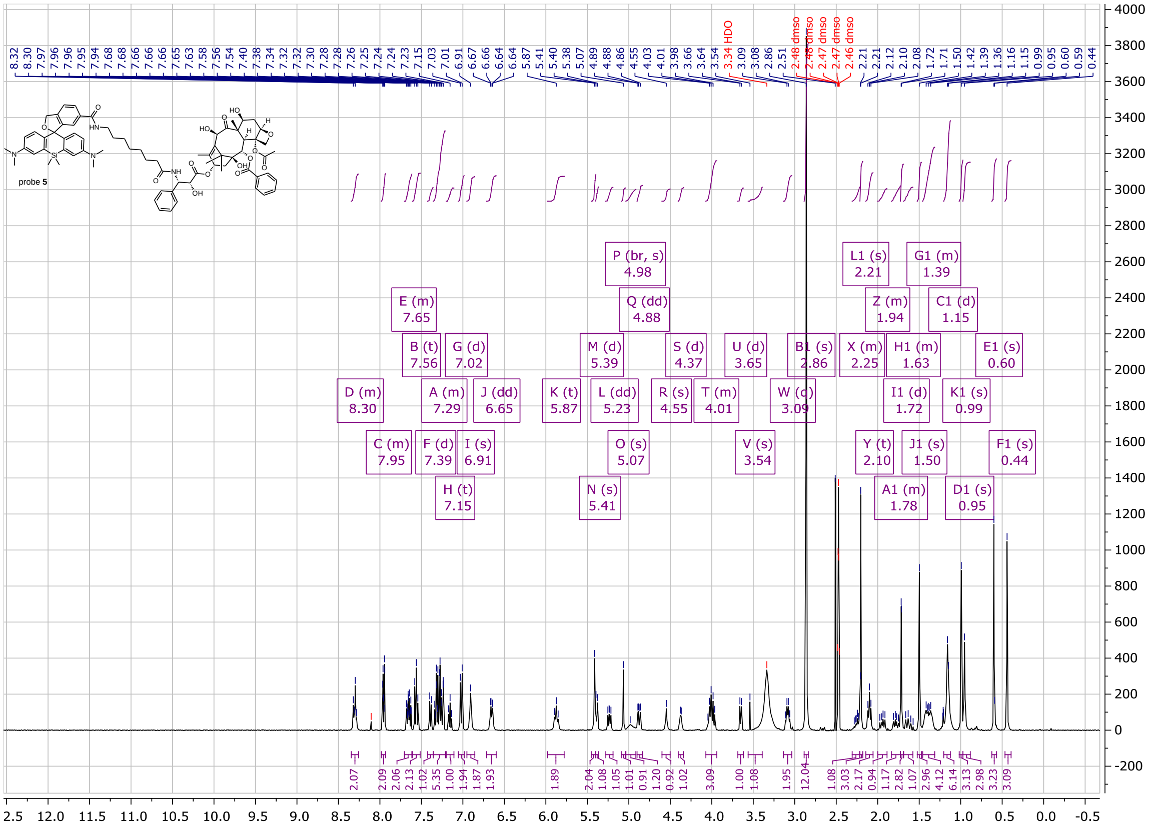

HRMS spectra copy of probe **5**

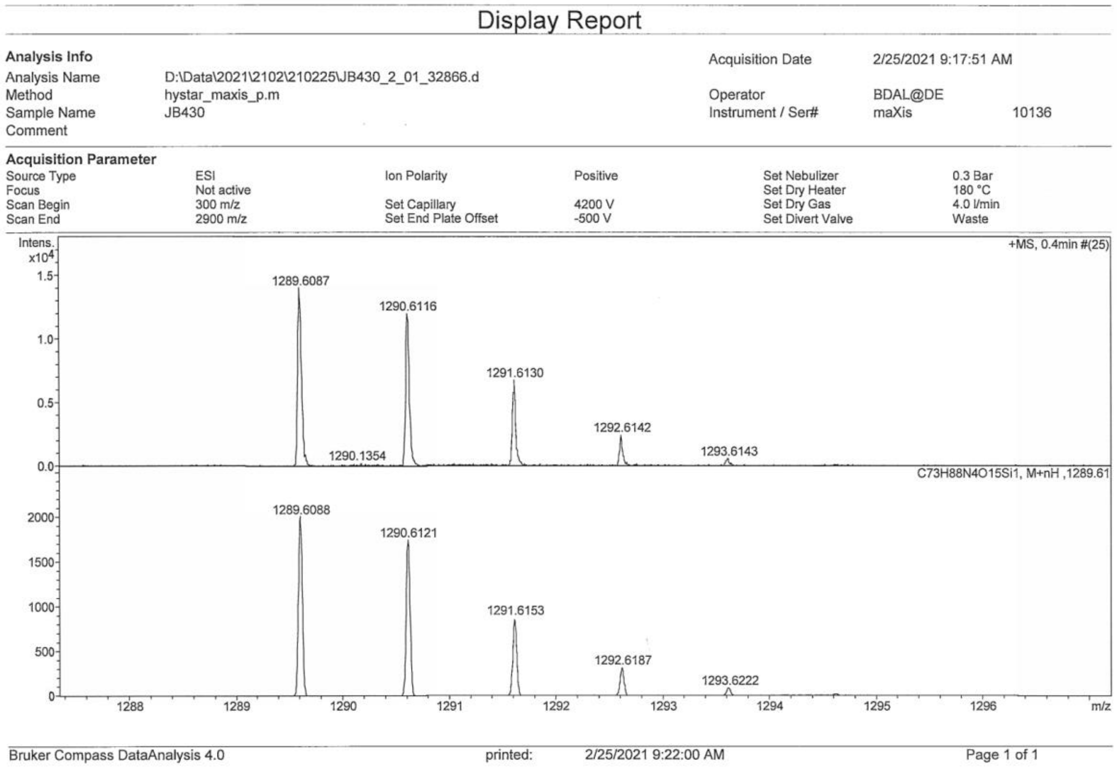

[1]H NMR spectra copy of probe **4**:

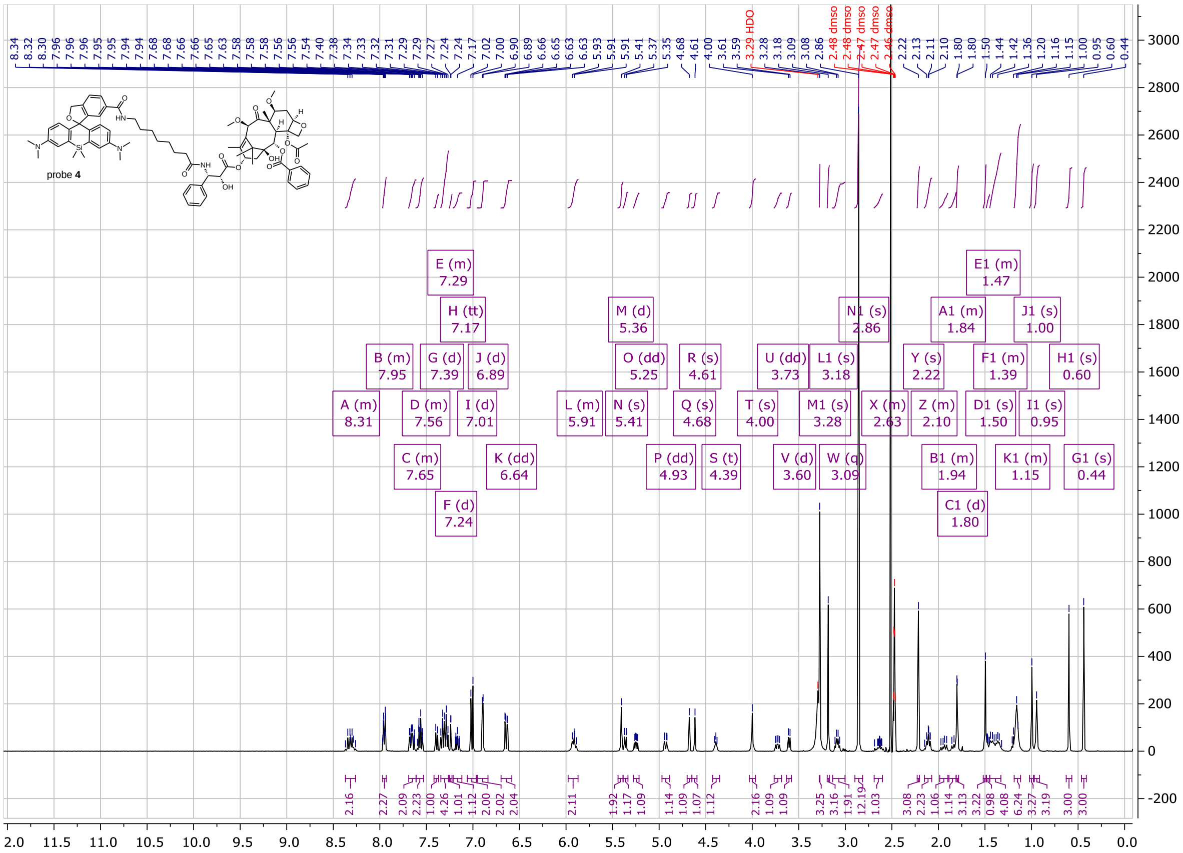

HRMS spectra copy of probe **4**:

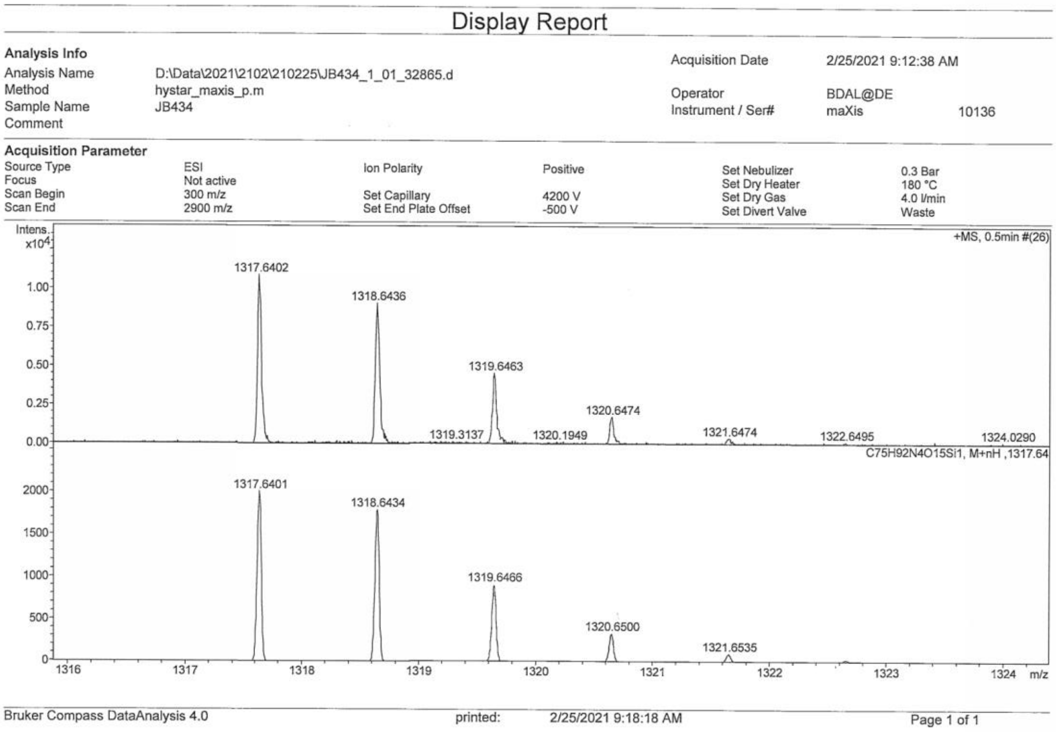

$^{1}$H NMR spectra copy of probe **3** (HMSiR-tubulin):

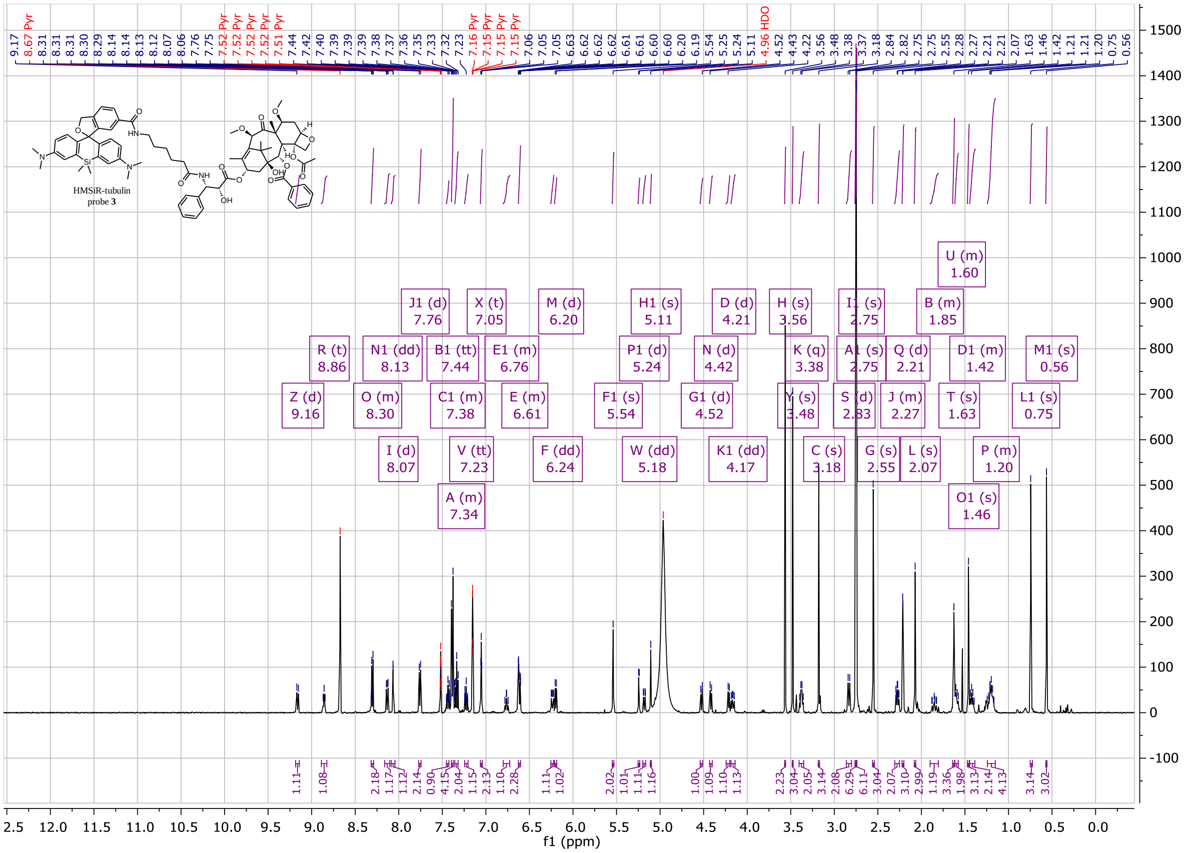

HRMS spectra copy of probe **3** (HMSiR-tubulin):

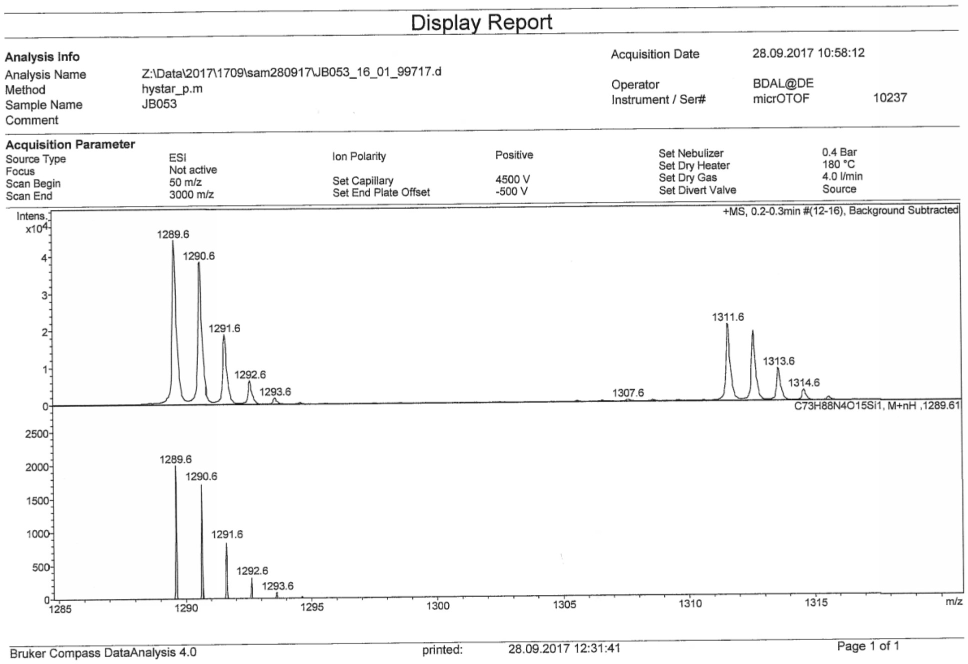

[1]H NMR spectra copy of probe **2**:

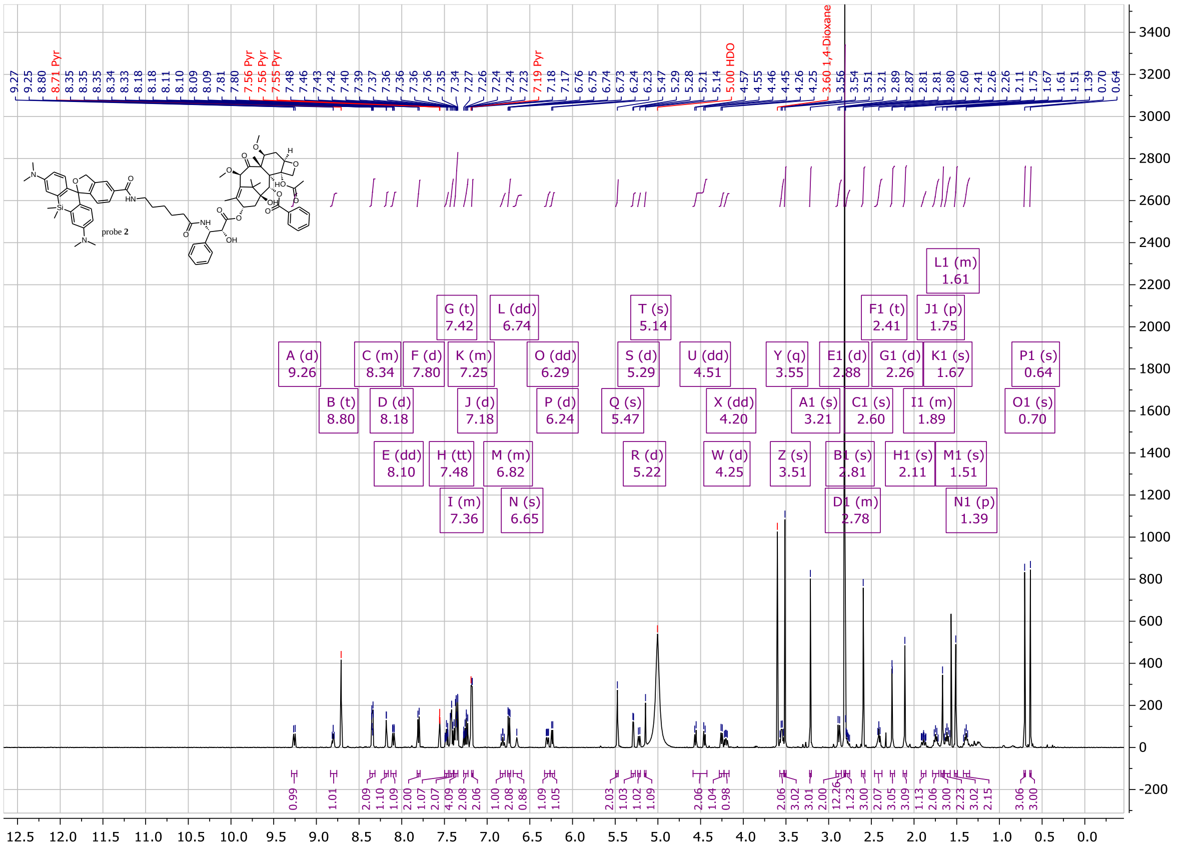

HRMS spectra copy of probe **2**:

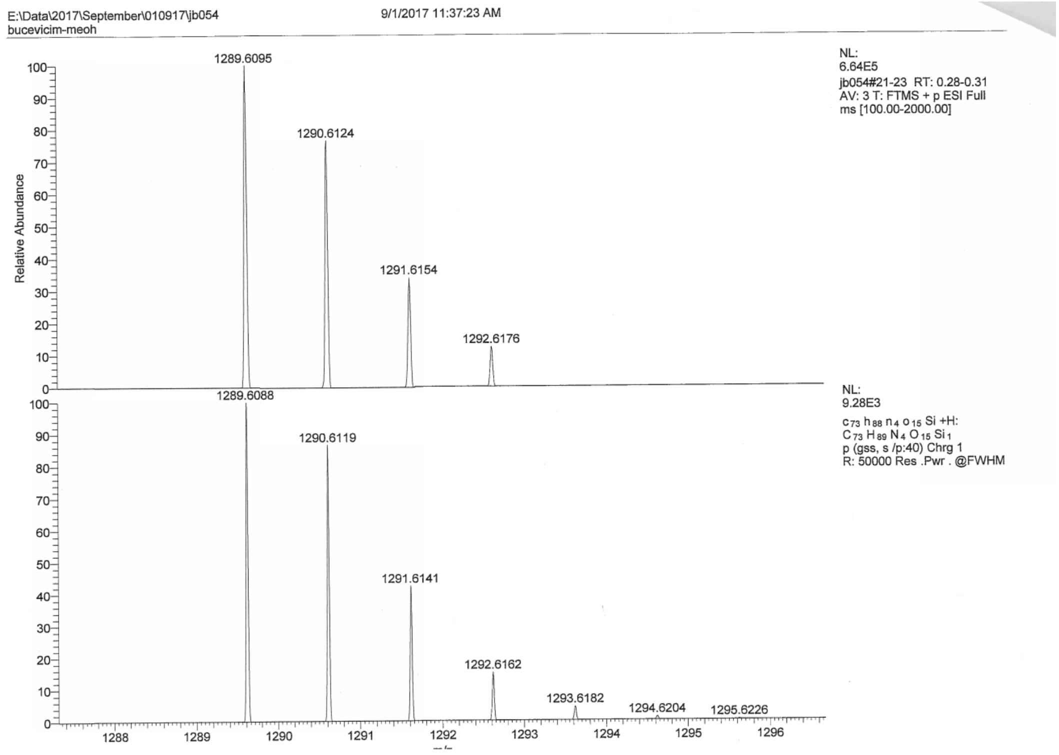

[1]H NMR spectra copy of probe **1**:

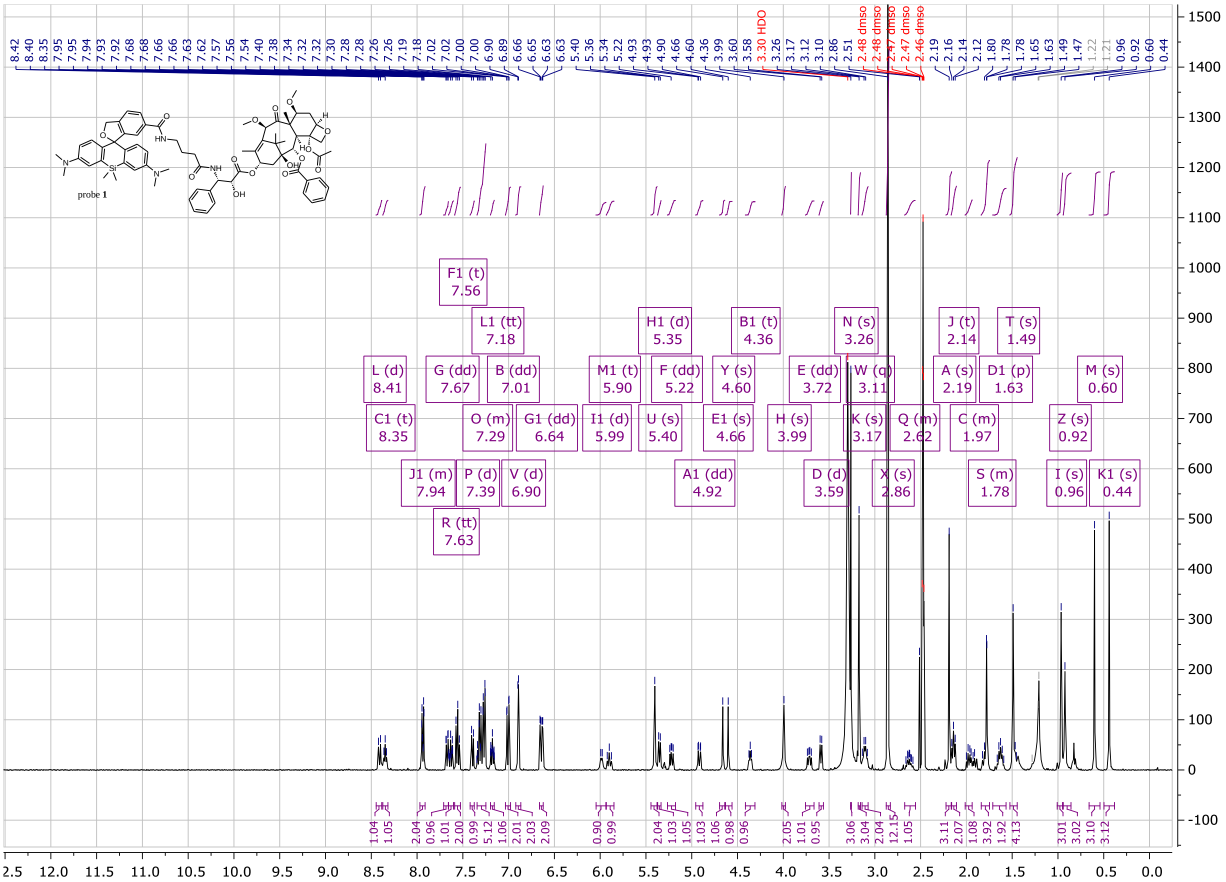

HRMS spectra copy of probe **1**:

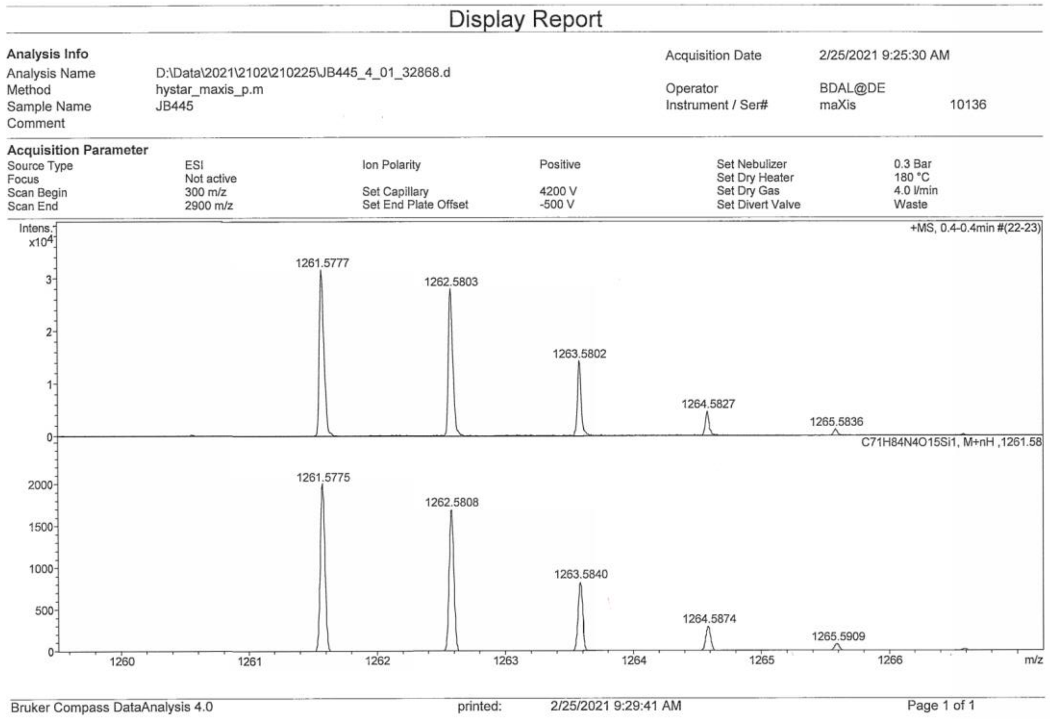

Display Report

| | | | |
|---|---|---|---|
| **Analysis Info** | | Acquisition Date | 2/25/2021 9:25:30 AM |
| Analysis Name | D:\Data\2021\2102\210225\JB445_4_01_32868.d | | |
| Method | hystar_maxis_p.m | Operator | BDAL@DE |
| Sample Name | JB445 | Instrument / Ser# | maXis 10136 |
| Comment | | | |

**Acquisition Parameter**

| | | | | | |
|---|---|---|---|---|---|
| Source Type | ESI | Ion Polarity | Positive | Set Nebulizer | 0.3 Bar |
| Focus | Not active | | | Set Dry Heater | 180 °C |
| Scan Begin | 300 m/z | Set Capillary | 4200 V | Set Dry Gas | 4.0 l/min |
| Scan End | 2900 m/z | Set End Plate Offset | -500 V | Set Divert Valve | Waste |

Bruker Compass DataAnalysis 4.0 printed: 2/25/2021 9:29:41 AM Page 1 of 1

$^{1}$H NMR spectra copy of probe **6**:

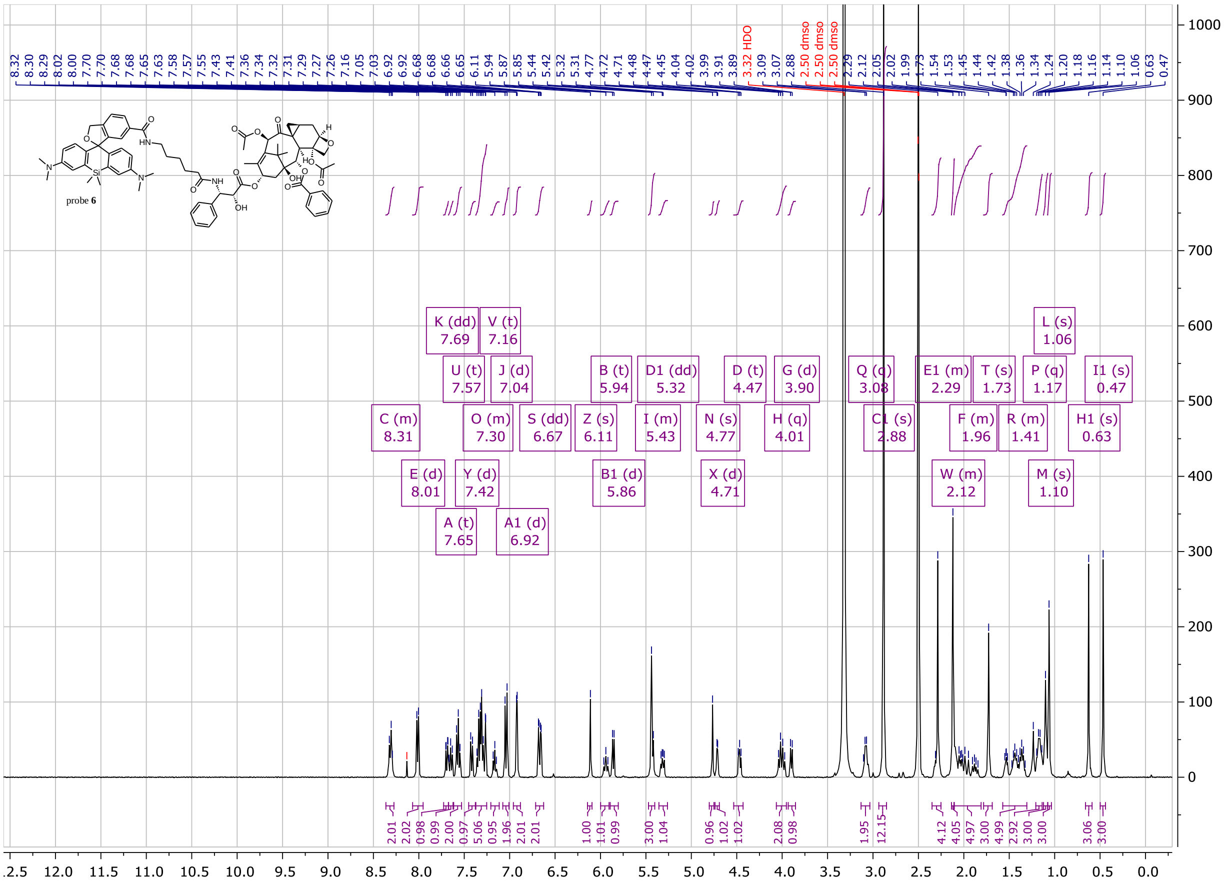

HRMS spectra copy of probe **6**:

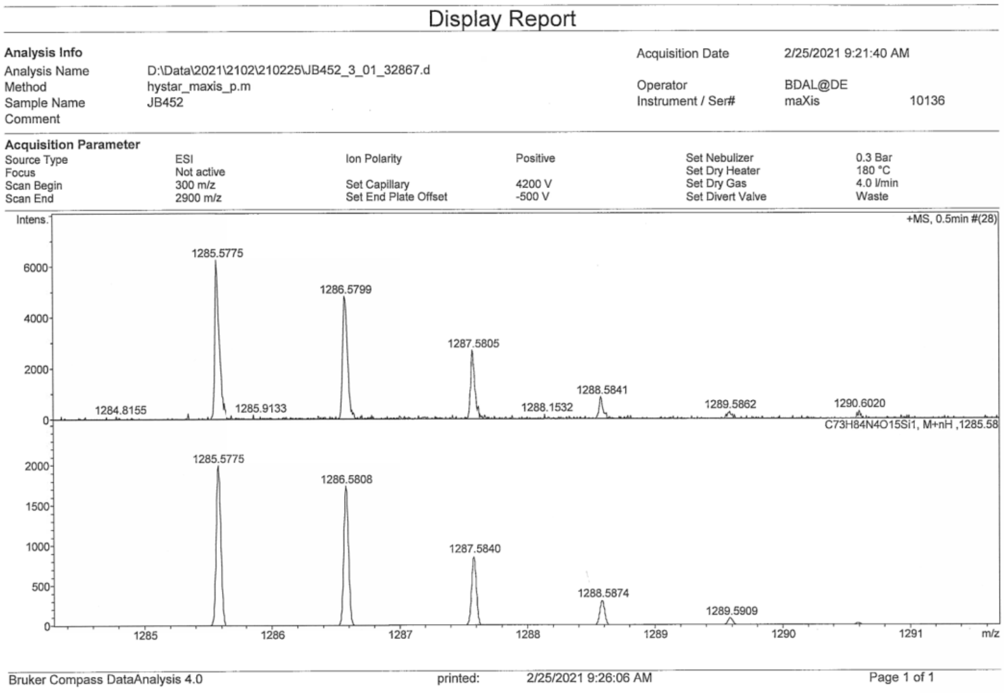

### References

1. Lukinavicius, G.; Mitronova, G. Y.; Schnorrenberg, S.; Butkevich, A. N.; Barthel, H.; Belov, V. N.; Hell, S. W., Fluorescent dyes and probes for super-resolution microscopy of microtubules and tracheoles in living cells and tissues. *Chem Sci* **2018,** *9* (13), 3324-3334.

2. Diaz, J. F.; Barasoain, I.; Andreu, J. M., Fast kinetics of Taxol binding to microtubules. Effects of solution variables and microtubule-associated proteins. *J Biol Chem* **2003,** *278* (10), 8407-19.

3. Diaz, J. F.; Strobe, R.; Engelborghs, Y.; Souto, A. A.; Andreu, J. M., Molecular recognition of taxol by microtubules. Kinetics and thermodynamics of binding of fluorescent taxol derivatives to an exposed site. *J. Biol. Chem.* **2000,** *275* (34), 26265-76.

4. D'Este, E.; Kamin, D.; Gottfert, F.; El-Hady, A.; Hell, S. W., STED nanoscopy reveals the ubiquity of subcortical cytoskeleton periodicity in living neurons. *Cell Rep* **2015,** *10* (8), 1246-51.

5. Schindelin, J.; Arganda-Carreras, I.; Frise, E.; Kaynig, V.; Longair, M.; Pietzsch, T.; Preibisch, S.; Rueden, C.; Saalfeld, S.; Schmid, B.; Tinevez, J. Y.; White, D. J.; Hartenstein, V.; Eliceiri, K.; Tomancak, P.; Cardona, A., Fiji: an open-source platform for biological-image analysis. *Nat. Methods* **2012,** *9* (7), 676-82.

6. Ovesny, M.; Krizek, P.; Borkovec, J.; Svindrych, Z.; Hagen, G. M., ThunderSTORM: a comprehensive ImageJ plug-in for PALM and STORM data analysis and super-resolution imaging. *Bioinformatics* **2014,** *30* (16), 2389-90.

7. Gottlieb, H. E.; Kotlyar, V.; Nudelman, A., NMR Chemical Shifts of Common Laboratory Solvents as Trace Impurities. *J Org Chem* **1997,** *62* (21), 7512-7515.

8. Ren, S.; Wang, Y.; Wang, J.; Gao, D.; Zhang, M.; Ding, N.; Li, Y., Synthesis and biological evaluation of novel larotaxel analogues. *Eur J Med Chem* **2018,** *156*, 692-710.

9. Bucevicius, J.; Gilat, T.; Lukinavicius, G., Far-red switching DNA probes for live cell nanoscopy. *Chem Commun (Camb)* **2020,** *56* (94), 14797-14800.
